# *Cladosporium* species detoxify multiple water micropollutants of emerging concern using diverse strategies

**DOI:** 10.1101/2023.09.25.559271

**Authors:** Maria Louise Leth, Kai Tang, Trine Sørensen, Aaron John Andersen, Claus Hélix-Nielsen, Birgitte Andersen, Jens Frisvad, Teis Esben Sondergaard, Henrik Rasmus Andersen, Maher Abou Hachem

## Abstract

The accumulation of micropollutants of emerging concern in aqueous systems raises safety concerns regarding biological systems and human health. Mycoremediation is a promising and green strategy to mitigate the micropollutant challenge. Hitherto, focus has mainly been on white-rot Basidiomycota and micropollutant transformation by ascomycetes remains underexplored. Here, we assayed 53 Ascomycota isolates from 10 genera for the removal of 22 micropollutants. Notably, 9 out of 22 micropollutants were removed from fungal culture supernatant at efficacies >45%. Temporal analysis of the nine top-performing strains, highlighted remarkable potency of *Cladosporium* isolates in removal of multiple micropollutants. Importantly, *Cladosporium* considerably reduced the toxicity of a micropollutant cocktail based on growth assays. Metabolomics analyses identified oxidation for 5-methyl-1H-benzotriazole and citalopram, whereas methylation and carboxylation were observed for 5-chlorobenzotriazole. No transformation products were detected for ciprofloxacin, sulfamethoxazole, and sertraline, hinting their extensive degradation. These findings suggest micropollutant transformation via diverse catalytic routes by *Cladosporium*. Genome sequencing and proteomic analyses of the top-performing isolates were consistent with the observed transformations and tentatively identified the molecular apparatus, conferring micropollutant transformation. This unprecedented study brings novel insight into the micropollutant transformation and detoxification capabilities of the prevalent *Cladosporium* species, thereby revealing a considerable and hitherto underappreciated potential of this genus and potentially other ascomycetes in micropollutant transformation.

**Importance:** At present, conventional wastewater treatment plants (WWTPs) are not designed for removing micropollutants, which are released into aqueous systems. This raises concerns due to the poor insight into micropollutant long-term interplay with biological systems. Innovating biotechnological solutions to tackle micropollutant require addressing the paucity of knowledge on microbial groups and molecular pathways, which mediate micropollutant transformation. Our study highlights the considerable potential of the *Cladosporium* genus that remains underexplored in the arena of micropollutant transformation. We report the first genomes sequences for three *Cladosporium* species: *C. allicinum, C. inversicolor,* and *C. fusiforme*, which sets the stage for further analyses of micropollutant transformation, but also offers an important resource on this ecologically significant, albeit under-studied genus and related Ascomycota.

## Introduction

The presence of micropollutants in aqueous systems, *e.g.* surface water, groundwater, and effluents from wastewater treatment plants (WWTPs) is an intensifying global challenge(1, 2). Micropollutants, which are generated by agricultural and industrial activities, as well as the widespread use of personal-care products and pharmaceuticals, are typically present in low concentrations (ng L^-1^-μg L^-1^) in wastewater(3). The “micropollutants of emerging concern” label has been issued for several compounds, awing to their potentially negative impacts on aquatic and terrestrial ecosystems, as well as human health(4, 5). Adverse effects on aquatic organisms, such as endocrine disruption, developmental abnormalities, and reduced reproductivity, have been observed(6, 7). These issues, which are exacerbated through bioaccumulation of certain micropollutants in the food chain(8), leading to human exposure, have emphasized the need to develop strategies to mitigate their adverse impacts.

Micropollutants, including analgesics, antibiotics, and anti-inflammatory drugs, are typically resistant to biological degradation(9), consistent with their presence in effluents from conventional activated sludge- type wastewater treatment plants (WWTPs) (10). Applying post-treatment physicochemical methods (*e.g.* ozonation) for upgrading existing treatment processes in WWTPs, improves micropollutants’ removal efficacy, but suffers from higher energy-intensiveness, operation costs, and potential generation of toxic by-products(11, 12). Mycoremediation is a promising eco-friendly strategy to remove micropollutants from wastewater(13). Fungi were reported to remove micropollutants from hospital wastewater(14) and antibiotics in wastewater(15). Specifically, white-rot fungi from the Basidiomycota phylum have proven effective in micropollutant removal(16). For example, *Trametes versicolor* was shown to remove a plethora of micropollutants, inducing naproxen, carbamazepine, and ibuprofen(14, 17). By comparison, the corresponding capacity of ascomycetes remains underexplored, despite their dominance in WWTPs(18–20). Mainly, *Aspergillus*(21) and *Penicillium oxalicum*(22) have been demonstrated to remove micropollutants. However, there is a knowledge gap regarding micropollutant transformation strategies and the identity of key players within Ascomycota.

Here we evaluated the removal of 22 diverse micropollutants (Fig. S1), featuring on wastewater watch lists in several European countries(23), by a panel of Ascomycota isolates. By combining microbiology, metabolomics, genome sequencing and differential proteomics, we identified *Cladosporium* strains potent in biotransformation and detoxification of several micropollutants. Distinct chemical modifications of micropollutants were identified and proteomics tentatively shortlisted candidate enzymes that mediate these biotransformation.

## Results

### *Cladosporium* members remove micropollutants with highest efficacy amongst 10 Ascomycota genera

We focused on Ascomycota members, harnessing the IBT culture (>38,000 fungal isolates from diverse habitats) collection at the Department of Biotechnology and Biomedicine, Technical University of Denmark). We hypothesized that fungi collected from contaminated sites, *e.g.* polluted soil or wastewater plants, are more likely to transform micropollutants, than counterparts from niches with no exposure to contaminants. Fifty three Ascomycota strains (Table S1), isolated at two different sites in Denmark: Lynetten WWTP (Copenhagen, Denmark) and trichloroethylene (TCE) polluted soil (Vedbæk, Denmark), were screened to examine their action on our selected micropollutant panel (Table S2). These isolates included 10 genera (*Aspergillus, Penicillium, Cladosporium, Fusarium, Stachybotrys, Geotrichum, Phoma, Monilochaetes, Onychocola,* and *Epicoccum*) from four classes (*Eurotiomycetes, Dothideomycetes, Sordariomycetes,* and *Saccharomycetes*) (Fig. 1A).

**FIG 1:**
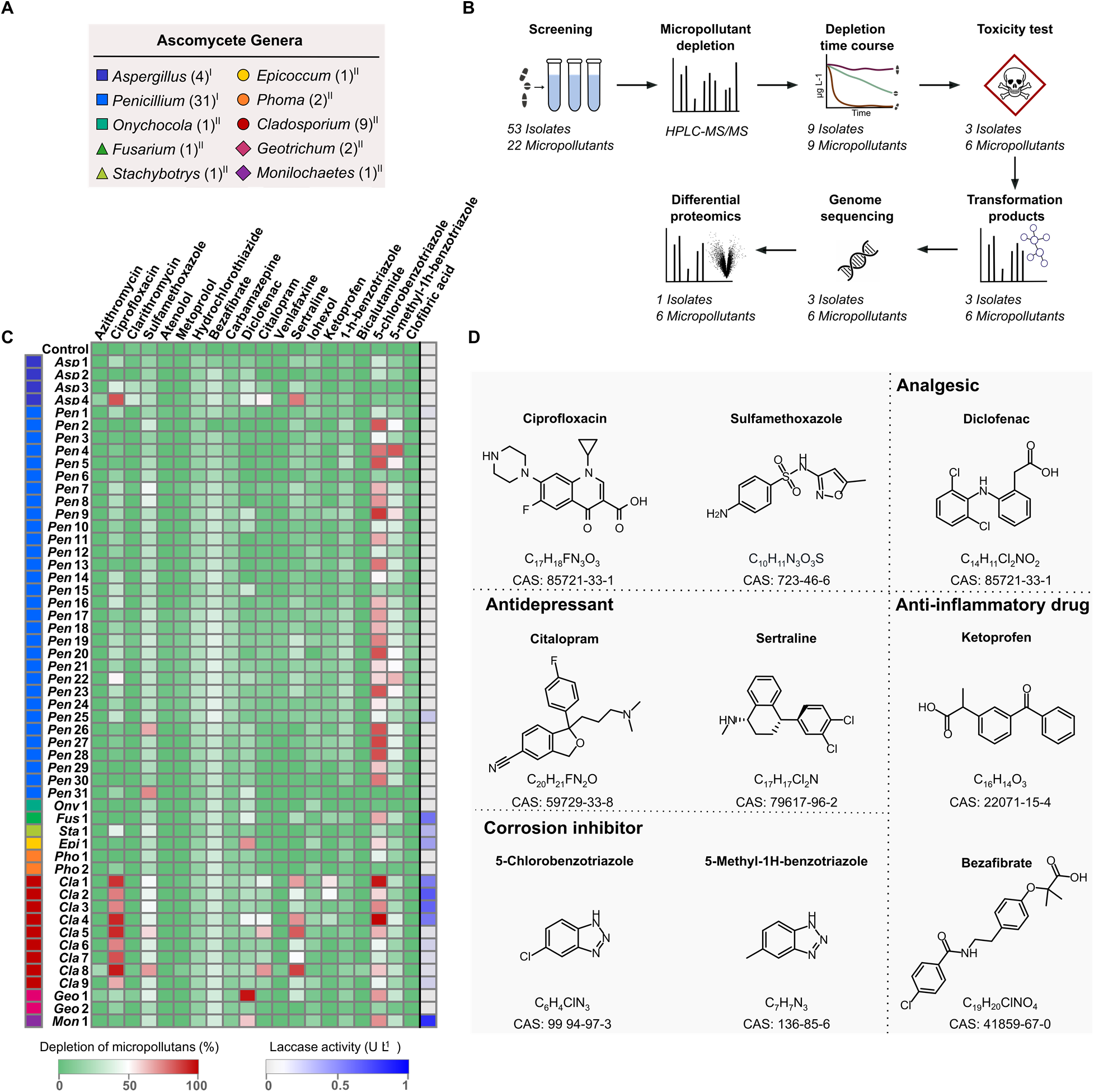
Micropollutant removal by the fungal isolate panel. A) The genus distribution of Ascomycota isolates in the present study and the number of isolates is in parentheses. The axonomic class is depicted as a symbol: Eurotiomycetes (square), Sordariomycetes (triangle), Dothideomycetes (circle), Saccharomycetes (diamond). ^I^Isolated from a wastewater treatment plant, ^II^Isolated from trichloroethylene (TCE) polluted soil. B) The Ascomycota micropollutant removal and analysis platform developed in this work. C) Heat map of micropollutant removal and laccase-like activities by the Ascomycota panel measured after 12 days incubation in minimal medium. *Asp: Aspergillus, Pen: Penicillium*, *Cla: Cladosporium, Fus: Fusarium, Sta: Stachybotrys, Geo: Geotrichum, Pho: Phoma, Mon: Monilochaetes, Ony: Onychocola,* and *Epi: Epicoccum.* The results are shown in percentage relative to an abiotic control. The results are from three independent biological replicates (n=3). D) Micropollutants that were removed ≥45% after 12 days incubation.

Micropollutants selections was based on a) recalcitrance to biotransformation, b) inclusion in Danish or EU watch lists, and c) structural/physiochemical diversity. This panel included pharmaceuticals (antibiotic, analgesics, antidepressants, anti-inflammatories, blood pressure regulators, lipid lowering agents, and X- ray contrast agents, besides corrosion inhibitors and herbicides (Fig. S1). Strain were screened in 6 mL cultures containing all micropollutants at 1 mg L^-1^ of each compound. A minimal medium (MM) and a nutrient-rich medium were used to investigate whether nutrient availability impacted micropollutant removal, based on monitoring micropollutant concentrations before and after 12 days of incubation using high-performance liquid chromatography-coupled triple quad mass spectrometry (HPLC-MS/MS) (Fig. 1B). The screening experiment in MM demonstrated growth of 50 out of 53 strains (94%) after 12 days and the removal of >45% of 9 out of 22 (41%) micropollutants, as compared to abiotic controls (Fig. 1C). Ciprofloxacin, diclofenac, and 5-chlorobenzotriazole were removed >95% by distinct strains. Overall, several genera removed certain micropollutants, but the most prominent was *Cladosporium*. The abiotic controls showed high removal of mefenamic acid (52%) and erythromycin (88%), suggesting this was an artefact, and these two compounds were excluded from further analyses (Table S3). We also measured secreted laccase activity (in culture supernatants), which was previously shown relevant for biotransformation of certain micropollutants(24). Laccase-like activity was detected for 8 out of 53 strains in MM (Fig. 1C), and appeared common in *Cladosporium* supernatants, but was also measured for *Fusarium, Stachybotrys, Monilochaetes,* and *Epicoccum* single isolates. Interestingly, the nutrient rich media sustained better growth of all 53 strains, however, poorer micropollutant removal was observed (Fig. S2, Table S4). Higher efficacy in MM, as compared to the nutrient-rich medium, indicated that the molecular machinery for the removal of micropollutants was upregulated during nutritional starvation.

To evaluate the presence of micropollutant transforming enzymes in the fungal secretomes, we analysed cell-free culture supernatants from day 6 and 10 with. No apparent removal of micropollutant was observed (Table S5), suggesting intracellular (or cell-surface attached) transformation systems.

### Cladosporium members remove multiple micropollutants with high relative efficacy

To analyse the kinetics of micropollutant removal, we selected the nine top-performing strains: *Penicillium* (*Pen*4, *Pen*9, *Pen*22)*, Aspergillus (Asp*4)*, Cladosporium,* (*Cla*4, *Cla*5, *Cla*8), *Geotrichum (Geo*1), and *Monilochaetes* (*Mon*1) (Fig. S3). These strains were grown for 15 days in 50 mL MM with a mixture of 9 micropollutants that were transformed at efficacies >45% in the screening experiment (Fig. 1D). The non-steroidal anti-inflammatory drugs diclofenac and ketoprofen were not efficiently removed within 15 days (Fig. 2A,B), with *Asp*4 mediating, being the top performer (63% removal of diclofenac). Additionally, *Asp*4 successfully removed >50% of the corrosion inhibitor 5-chlorobenzotriazole (Fig. 2C), while the removal of the similar corrosion inhibitor 5-methyl-1-h-benzotriazole was <25% (Fig. 2D). The concentration of the antibiotic ciprofloxacin already decreased by 50% in the *Asp*4 culture, as compared to the control at day 0 and day 3, but increased hereafter, suggesting initial sorption to the fungal biomass, followed by partial release into the medium (Fig. 2E). Notably, the three penicillia exhibited different capabilities. The removal of micropollutants was <30% by *Pen*9 and *Pen*2, except for of the anti-inflammatory drug bezafibrate, which was removed >70% (Fig. 2F). By contrast, *Pen*4 removed multiple micropollutants, including >90% of the two corrosion inhibitors, 5-methyl-1H-benzotriazole and 5-chlorobenzotriazole (Fig. 2C,D). Interestingly, *Pen*4 removed 5-methyl-1H-benzotriazole completely by day 6, and >60% of the antibiotic sulfamethoxazole was also removed within this timeframe (Fig. 2G). One of the more efficient strains was *Mon*1, which removed >90% of ciprofloxacin, sertraline, and citalopram (Fig. 2E, H, I) and >40% of 5-chlorobenzotriazole, 5-methyl-1H-benzotriazole, and sulfamethoxazole (Fig. 2C,D,G). Based on the initial screening, we selected three *Cladosporium* strains (*Cla*4, *Cla*5, and *Cla*8) for the time course analysis. These strains removed >95% of ciprofloxacin, sertraline, and citalopram within 15 days (Fig. 2E,G,H), whereas 25-90% of 5-chlorobenzotriazole, 5-methyl-1H-benzotriazole, and sulfamethoxazole were also removed (Fig. 2C,B,G). The top performing strain was *Cla*4 based on removal efficacies >90% for six out of the nine tested micropollutants.

**FIG. 2:**
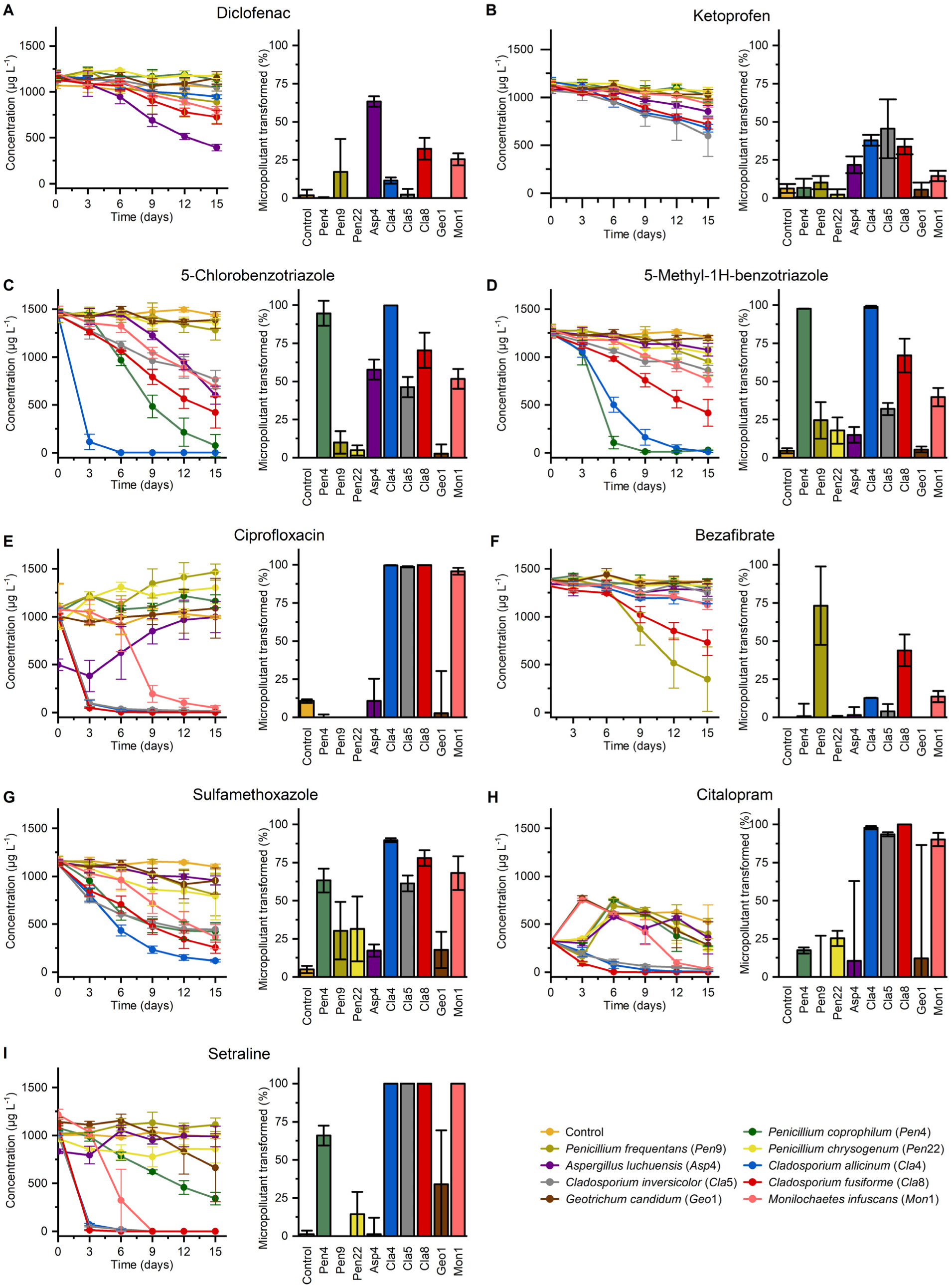
Time course analysis of micropollutant removal by selected Ascomycota strains. The micropollutants were in a mixture with a concentration of 1 mg L^-1^ of each compound and the growth was in minimal media for 15 days at room temperature. A-I represent the removal time course of the micropollutants; diclofenac, ketoprofen, 5-chlorobenzotriazole, 5-methyl-1H- benzotriazole, ciprofloxacin, bezafibrate, sulfamethoxazole, citalopram, and sertraline, respectively. Data are shown as an average of a biological triplicate (n=3) with standard deviations.

### Cladosporium isolates mediate the detoxification of multiple micropollutants

The three *Cladosporium* isolates *Cla*4, *Cla*5 and *Cla*8 mediated efficient removal of six micropollutants (5- methyl-1H-benzotriazole, 5-chlorobenzotriazole, ciprofloxacin, sulfamethoxazole, sertraline, and citalopram) as shown above. The observed removal may stem from physical biosorption of the micropollutant to the microbial biomass (*e.g.* the mycelium surface) or enzymatic transformation. To evaluate this, we analysed sorption to *Cla*4, *Cla*5, and *Cla*8 by extraction and quantification of the micropollutants from fungal biomass after 15 days and comparison to an abiotic control. The contribution of sorption was negligible for the micropollutants, except for 38% and 26% for the two anti-depressants sertraline and citalopram, respectively, onto *Cla*4 (Fig. 3A). These results strongly indicated that biotransformation was mainly behind the observed micropollutant removal. A similar trend was observed for *Cla*5, however, the sorption extent to the anti-depressants were lower (15% and 11%). The data from *Cla*8 were less compelling due to a high level of sorption for 5-methyl-1H-benzotriazole, 5- chlorobenzotriazole, and sulfamethoxazole in addition to the anti-depressants, although we cannot exclude that internalization of the compounds by the isolates could also contribute to the observed sorption.

**FIG 3:**
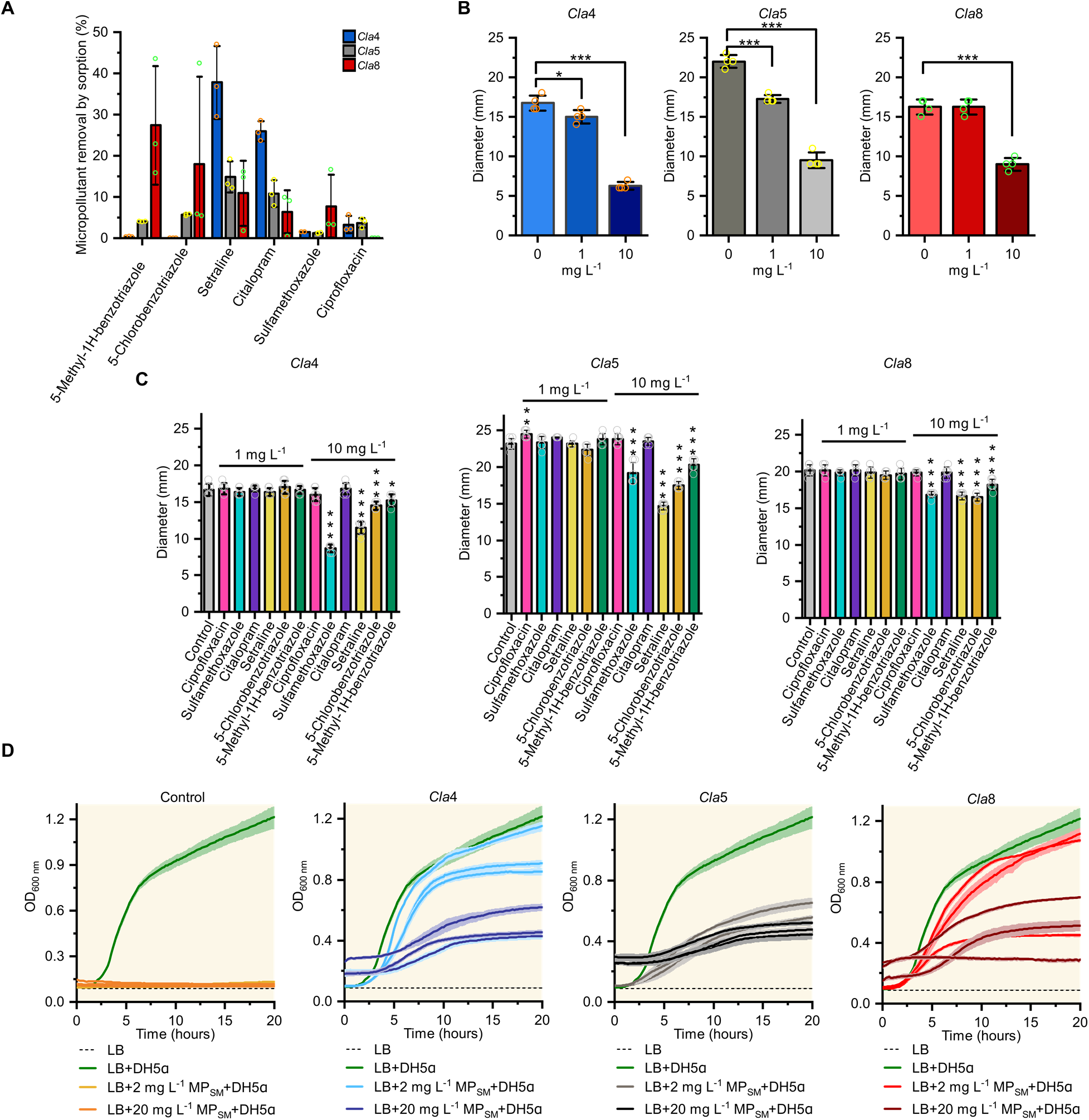
The detoxification of micropollutants by *Cladosporium* isolates (*Cla*4*, Cla*5 *and Cla*8). A) Sorption of micropollutant to fungal biomass after 15 days. The micropollutants were in a mixture with an initial concentration of 1 mg L^-1^ of each compound. The results are shown relative to an abiotic control and plotted as mean from an independent biological triplicate with standard deviations. B-C) Inhibition of fungal growth by a micropollutant mixture or single compounds, respectively. The fungi were grown on MM agar plates supplemented with the micropollutants 5-methyl-1H-benzotriazole, 5-chlorobenzotriazole, ciprofloxacin, sulfamethoxazole, sertraline, and citalopram. The concentration of each micropollutant in the mixtures and on the plates with a single compound was 0 mg L^-1^, 1 mg L^-1^, or 10 mg L^-1^. Results are shown as an average of the diameter from four colonies (n=4). Statistical significance of the comparisons to the controls was calculated with two sample t-test: *, *p*-value < 0.05; **, *p*-value < 0.01; ***, *p*-value < 0.001. D) Detoxification of the same mixture of micropollutants in panel C after incubation with the *Cladosporium* isolates for 15 days at a micropollutant concentration of 1 mg L^-1^. The detoxification was assayed by lyophilisation of the spent medium (SM) at the end of the incubation with the fungi and reconstituting it to 2 mg L^-1^ or 20 mg L^-1^ of each individual micropollutant (MP) in LB medium, which is used for growth of *E.coli* DHα. The data are from three biological replicates each performed with a technical triplicate for *E.coli* DHα growth. Coloured shaded regions around the growth curves (mean) indicate the standard deviations of the technical replicate (n=3).

Next, we assayed the toxicity of a mixture of the six efficiently-removed micropollutant against the same strains. This assay, which was based on growth inhibition (changes in fungal colony diameter) on MM plates, revealed dose dependent toxicity (Fig. 3B). To rank the toxicity of individual components, growth in the presence of each micropollutants was performed (Fig. 3C). Compared to a control without micropollutants, the diameter of the fungal colonies decreased substantially on 5-methyl-1H- benzotriazole, 5-chlorobenzotriazole, sulfamethoxazole, and sertraline for all three strains, indicating that these micropollutants were the most toxic in the mixture. Interestingly, sulfamethoxazole was most toxic for *Cla*4, which was the most efficient in transforming this antibiotic (Fig. 2D). The correlation between toxicity and removal efficacy, suggested that the removal maybe an evolutionary adaptation to detoxify similar natural compounds or synthetic xenobiotic analogues.

To evaluate whether the incubation with the fungal strains impacts toxicity, we used a common assay based on *E. coli* DH5α growth inhibition on the selected micropollutants before and after exposure to the fungal strains. The assay, rapidly provides semi-quantitative toxicity data, as compared to the above analysis. The minimum inhibitory concentration of the two antibiotics ciprofloxacin and sulfamethoxazole was reported to be 0.02 and 1.2 mg L^-1^, respectively for *E. coli*(25), which was consistent with abolished growth in the micropollutant-containing medium prior fungal incubation (Fig. 3D). Strikingly, the supernatants from the fungal cultures *Cla*4 and *Cla*8 with the same micropollutants at 2 mg L^-1^ (corresponding to the initial concentration) failed to inhibit *E. coli* DH5α, which grew comparably to the LB medium control. Even the 10-fold higher (20 mg L^-1^) concentration supported growth of *E. coli* DH5α after individual treatment with the three isolates. The results provide compelling evidence that the fungal isolates mediated efficient detoxification of the micropollutant mixture.

### Metabolomics analyses reveal different transformation routes of micropollutants

To bring insight into the fate of the transformed micropollutants, supernatants of the top micropollutant- transforming strains *Cla*4, *Cla*5, and *Cla*8 grown on a mixture of the six micropollutants mixture above were analysed using LC-MS/MS combined with non-targeted molecular network analysis. This analysis allows for identification of minor chemical modifications, via detection of related emerging species of similar fragmentation patterns as the un-modified micropollutants. For *Cla*4, oxidation of 5-methyl-1H- benzotriazole by the incorporation of either a single oxygen atom (P1) or two oxygen atoms (P2) in this compound was demonstrated (Fig. 4A). The initial oxidation product, P1, was detected at day 6, but was not detected by day 15. Conversely, the second oxidation product, P2, exhibited an increasing trend over time, indicating sequential double oxidation. By contrast, *Cla*5 and *Cla*8, which showed lower efficacy in removal of this micropollutant, primarily produced a single oxidation product, P1, after the 15-day period, while the double-oxygenated product was not reliably detected (Fig. 4B,C). The molecular structures of two corrosion inhibitors, 5-methyl-1H-benzotriazole and 5-chlorobenzotriazole, are nearly identical, differing only in the presence of a methyl- and a chloro-group, respectively. Surprisingly, network analysis of the supernatant of *Cla*4 did not identify any transformation products of 5-chlorobenzotriazole, despite its complete removal within six days (Fig. 4A). *Cla*5 and *Cla*8 exhibited lower efficacy in removing this compound, however, two transformation products (P3 and P4), sharing the same molecular mass (m/z [M+H]+=226.0378), increased abundance from day 3 to day 15. This mass corresponds to 5- chlorobenzotriazole+C_3_H_4_O_2_, indicating a double methylation and addition of a carboxyl group. All three strains exhibited high efficacy in removing citalopram, with <5% remaining in the supernatants after 15 days. A single product (P5) was detected in all three supernatants, resulting from the oxidation of citalopram by the addition of two oxygen atoms. Remarkably, the abundance of P5 decreased in the *Cla*8 supernatant after day 3 and was barely detectable by day 15.

**FIG 4:**
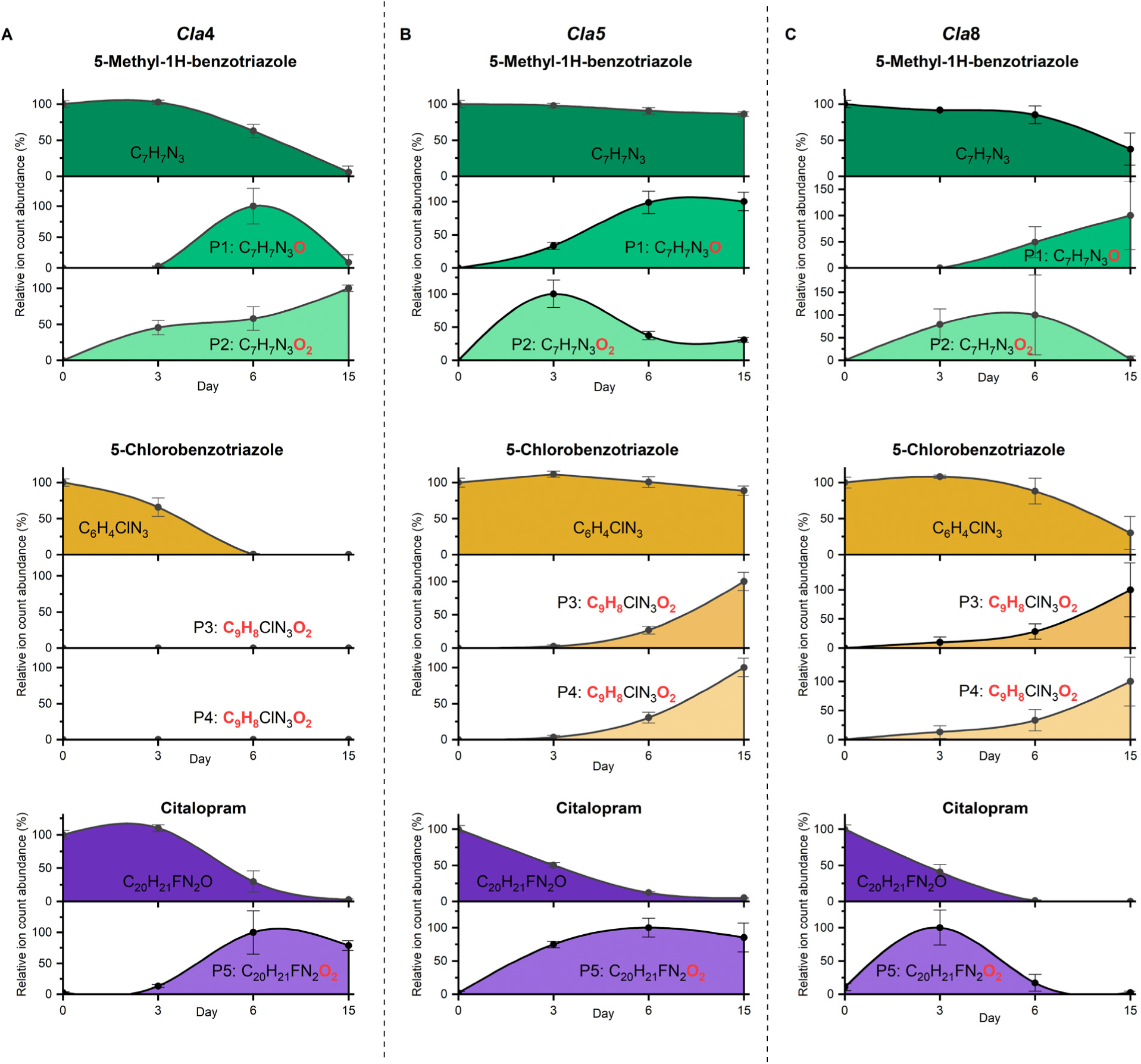
Metabolomics analysis of the transformation products of the micropollutants by *Cladosporium* isolates. Fungal supernatants of *Cla*4, *Cla*5, and *Cla*8 grown on a mixture of 5-methyl-1H-benzotriazole, 5-chlorobenzotriazole, ciprofloxacin, sulfamethoxazole, sertraline, and citalopram (1 mg L^-1^ of each micropollutant) were analysed using LC-MS/MS combined with non-targeted molecular network analysis. P3 and P4 were not detected for *Cla*4. The results are an average of independent experiments (n=4) with standard deviations relative to the individual features.

In summary, we identified two transformed products for the corrosion inhibitors 5-methyl-1H- benzotriazole (P1, P2) and 5-chlorobenzotriazole (P3, P4), as well as a single product for citalopram (P5). The observed changes in molecular mass strongly suggest that these transformations involve two levels of oxidation and methylation. Interestingly, our analysis did not reveal any transformed products for ciprofloxacin, sulfamethoxazole, and sertraline, which suggests that these compounds may have undergone more extensive/complete degradation.

### Genome sequencing and comparative genomic analyses

There was no available genomes for the three top performing Cladosporium, isolates were from soil contaminated with the industrial solvent trichloroethylene (TCE, C_2_HCl_3_). In fact, only 23 genomes from the *Cladosporium* genus are available from NCBI to date. Therefore, we sequenced the three strains to allow for detailed analyses on the molecular apparatus that confers the observed biotransformations.

Long read sequencing generated 2064373344, 3673242697, and 1937775533 bases across 174766, 459240, and 344131 reads for *Cla*4, *Cla*5, and *Cla*8, respectively. After quality filtering 1658648579, 2470711051, and 1031342831 bases and 66629, 105175, and 49906 reads were used for *de novo* assembly, which resulted in three draft genomes of 30.7, 33.9 and 33.7 Mb for *Cla*4, *Cla*5, and *Cla*8 genomes, respectively (Table S7). Contig length N50 and N99 together with Benchmarking Universal Single-Copy Orthologues (BUSCO) metrics showed established the high quality of the assembled genomes. Annotation of the genomes resulted in the identification of 11515, 12685, and 11651 protein-coding genes of which 8949 (77.7%), 9582 (75.5%), and 9119 (78.9%), respectively, could be assigned a function (Table S7).

A BLASTn analysis of the internal transcribed spacer regions (ITS) of the rDNA, partial fragments of actin (*act*) and the translation elongation factor 1-α (*tef1*) gene loci(26) assigned the genomes of *Cla*4, *Cla*5, and *Cla*8 as *Cladosporium allicinum, Cladosporium inversicolor,* and *Cladosporium fusiforme* species, respectively (Table S8). Henceforth these strains will be referred to as *C. allicinum* IBT 42152*, C. inversicolor* IBT 42153, and *C. fusiforme* IBT 42164. Three major complexes of species are recognized within *Cladosporium* and used for taxonomic analyses, *viz.* the *C. herbarum, C. sphaerospermum* and *C. cladosporioides* species complexes(27). *C. allicinum, C. inversicolor,* and *C. fusiforme* belong to the *C. herbarum, C. sphaerospermum* and *C. cladosporioides,* complexes, respectively. Pairwise whole genome comparisons using genomic average nucleotide identity (ANI), demonstrated that the three isolates cluster with species from respective complex (based on their high ANI values) (Fig. S4). In conclusion, these genomes are the first reported for all three species, which provides an important resource for comparative genomic analyses, beyond this work.

To investigate the molecular basis for micropollutant conversion, we analysed the predicted cytochrome P450 repertoire of the sequenced *Cladosporium* isolates. This enzyme family harbours diverse haeme- thiolate oxidoreductases (mainly monooxygenases) that play crucial roles in primary and secondary metabolism, as well as in detoxification of xenobiotics such as environmental pollutants and plant-derived toxins(28). *C. allicinum* IBT 42152 (*Cla*4), *C. inversicolor* IBT 42153 (*Cla*5), and *C. fusiforme* (*Cla*8) encoded 82, 85, and 76 genes classified as P450, respectively, based on eggNOC (Fig. 5). While the majority of *Cladosporium* genomes possess similar P450 repertoires, distinct members showed about 50% reduction in the P450 pool, indicating genomic adaptations to different ecological niches, with respect to secondary metabolism including xenobiotic transformation. We also analysed the carbohydrate active enzyme arsenal (CAZyome) due to the potential of certain CAZy-assigned auxiliary activity (AA) oxidoreductases (29) to act on some xenobiotics, *e.g.* laccases (AA1) or flavoenzymes from the vanillyl alcohol oxidase (VAO; EC 1.1.3.38) superfamily.

**FIG 5:**
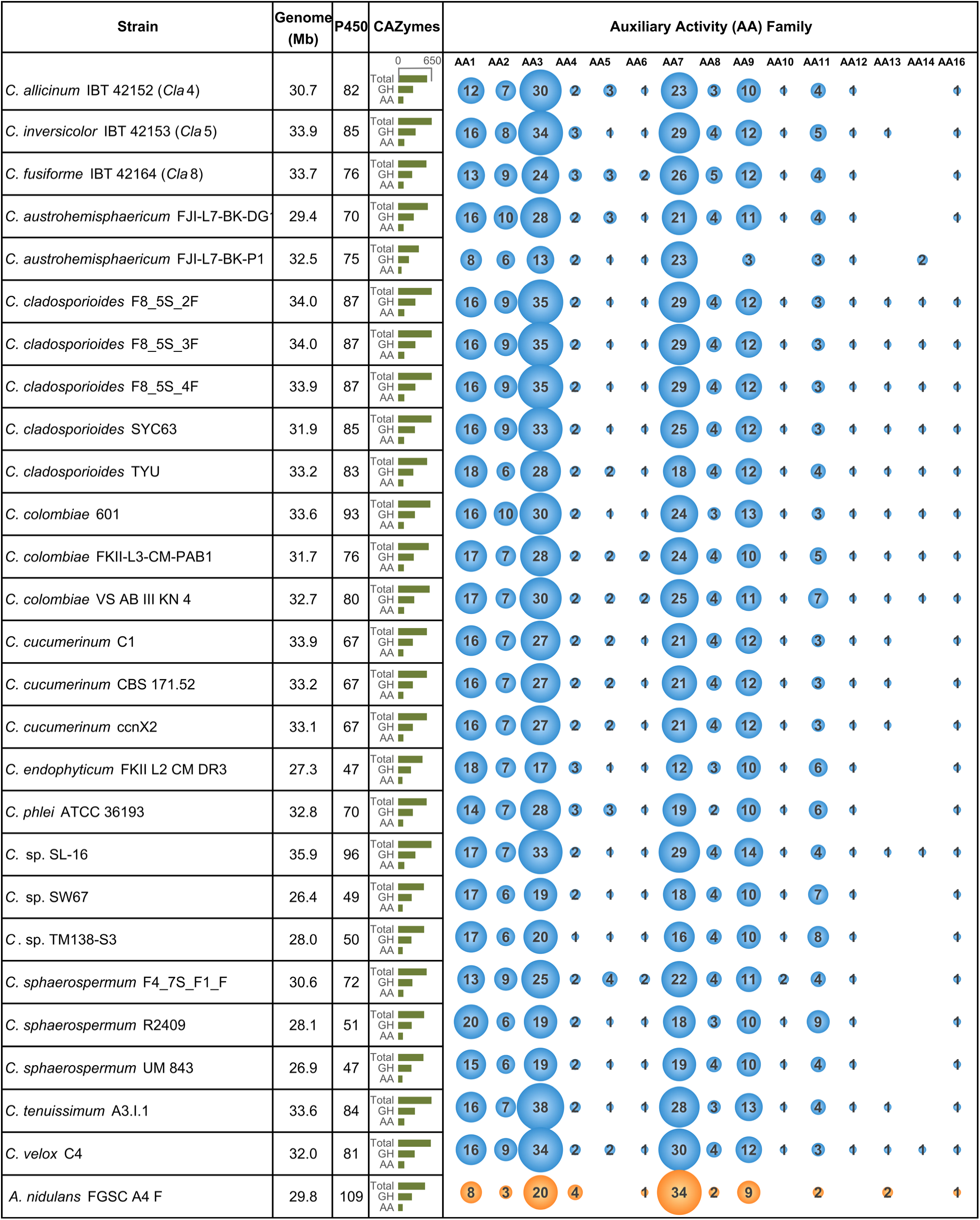
Comparative genomics of sequenced *Cladosporium* species*. Aspergillus nidulans* FGSC A4 F was included as outgroup. Colored circle shows the number of enzymes within each auxiliary activities (AA) family and the size of each circles reflects this. Circle colors; blue: *Cladosporium*, orange: *Aspergillus*.

Redox-active CAZymes from 15 AA families were identified. Although most AA enzymes are implicated in the degradation of lignocellulose. Laccases (AA1) and peroxidases (AA2) possess high redox potentials and broad specificities, qualifying them to act on non lignocellulosic substrates(30). Our analyses revealed generally similar distribution of AA families in the *Cladosporium* genomes, with an overrepresentation of flavo-enzymes from AA3 and AA7, both harbouring dehydrogenases and oxidases(31). Interestingly, the content of potential laccases (AA1) and peroxidases (AA2) was relatively higher in the *Cladosporium* genomes as compared to the model saprotroph *Aspergillus nidulans*. By contrast, several *Cladosporium* genomes lacked AA13 genes (starch-active lytic polysaccharide monooxygenase, LPMO) and other lignocellulose active LPMOs from AA14 (xylan-active) and AA16 (cellulose-active), which is valid for the three sequenced species. Reduction in LPMO genes may reflect adaptations to niches with alternative and more abundant metabolic resources, such as contaminated soils.

### Multiple oxidoreductases upregulated in the presence of micropollutants

To investigate the molecular apparatus upregulated during growth on micropollutants, we performed a differential proteomic analysis on the most potent isolate, *C. allicinum* IBT 42152 (*Cla*4), in presence and absence of the same six micropollutants as above. We analysed both the intracellular proteome and the secretome. Only 43 proteins were upregulated (log_2_≥ 1.5, *p-*value≤ 0.05) in the secretome (Table S10). However, only three of these proteins had predicted signal peptides, one of which was a predicted FAD-dependent monooxygenase (Q7P36_000751, Table S10). This, together with poor separation between the samples with micropollutants and the controls, raised concerns on the validity of this data, and we chose not to pursue this analysis (Table S10, 11).

The intracellular proteome suggested a stress response with more than 3-fold downregulated proteins as upregulated counterparts. Thus, 44 proteins were upregulated (log_2_≥ 1.5, *p-*value≤ 0.05), whereas 138 proteins were downregulated (log_2_≤ −1.5, *p*-value≤ 0.05) in presence of the micropollutants versus the control (Table 1, Table S12, 13). Clusters of Orthologous Groups analysis showed that the majority of the down-regulated proteins were grouped into the G category involving carbohydrate transport and metabolism (Fig. 6). In this group, 22 proteins were predicted to be glycoside hydrolases (GHs), mainly glucosidases and glucanases (Table S13). Glucans are the most abundant polysaccharides in the cell walls of fungi(32). The observed responses may reflect reduced cell-wall synthesis/remodelling.

**FIG 6:**
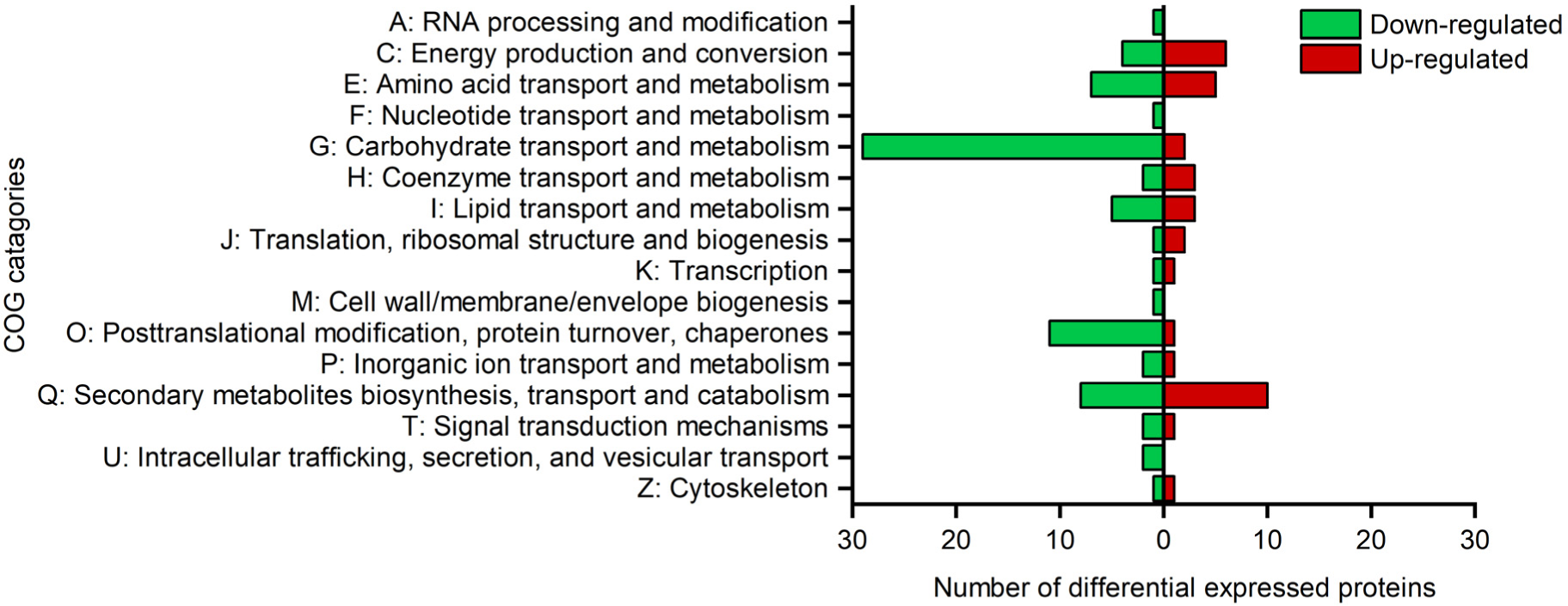
Functional annotation of intracellular differentially expressed proteins. Intracellular proteins with log_2_ fold change ≥ 1.5 and ≤ −1.5 were assigned into Clusters of Orthologous Groups (COG) categories by eggNOC-mapper. Proteins assigned >1 COG’s are represented in both categories. Proteins assigned into the “function unknown” category (7 up-regulated and 38 down- regulated) and are not shown in the figure.

**Table 1:** Differential upregulation of intracellular oxidoreductases and methyltransferases in the presence of micropollutants.

| Accession | Name | Log2FC | COG | SP | Localization |
| --- | --- | --- | --- | --- | --- |
| Q7P36_007294 | Aldo/keto reductase | 3.1 | C | no | Cytoplasm |
| Q7P36_010724 | Aldo/keto reductase | 3.0 | C | no | Cytoplasm |
| Q7P36_007278 | Methyltransferase | 2.9 | S | no | Cytoplasm, Nucleus |
| Q7P36_007637 | Short-chain dehydrogenase/reductase | 2.5 | Q | no | Cytoplasm |
| Q7P36_004137 | NAD- or NADP-dependent oxidoreductases | 2.4 | Q | no | Cytoplasm |
| Q7P36_006523 | Zinc-binding oxidoreductase | 2.4 | C | no | Cytoplasm |
| Q7P36_007820 | Baeyer-Villiger monooxygenase | 2.1 | Q | no | Peroxisome |
| Q7P36_001385 | Aldo/keto reductase | 2.0 | S | no | Cytoplasm |
| Q7P36_008506 | 2-Hydroxyacid oxidase | 1.9 | C | no | Peroxisome |
| Q7P36_010634 | Aldo/keto reductase | 1.9 | S | no | Cytoplasm, Nucleus |
| Q7P36_000482 | Short-chain dehydrogenase/reductase | 1.8 | Q | no | Cytoplasm |
| Q7P36_002133 | Sulfide quinone-reductase | 1.8 | C | no | Mitochondrion |
| Q7P36_008457 | Phosphatidylethanolamine *N*-methyltransferase | 1.7 | I | no | Endoplasmic reticulum |
| Q7P36_006997 | S-adenosyl-L-methionine-dependent methyltransferase | 1.7 | Q | no | Cytoplasm, Nucleus |
| Q7P36_008529 | Short-chain dehydrogenase/reductase | 1.7 | Q | no | Cytoplasm |
| Q7P36_004165 | 3-β-hydroxysteroid dehydrogenase/isomerase | 1.6 | E,I | no | Endoplasmic reticulum |
| Q7P36_010397 | Pyridine nucleotide-disulphide oxidoreductase | 1.6 | Q | no | Cytoplasm |
| Q7P36_001318 | S-adenosyl-L-methionine-dependent methyltransferase | 1.6 | S | no | Cytoplasm, Nucleus |
| Q7P36_002807 | Thioredoxin | 1.6 | O | no | Cytoplasm |
| Q7P36_008349 | Short-chain dehydrogenase/reductase | 1.6 | Q | no | Cytoplasm |
| Q7P36_000623 | Malate oxidoreductase | 1.5 | C | no | Mitochondrion |
| Q7P36_009238 | NAD- or NADP-dependent oxidoreductases | 1.5 | S | no | Cytoplasm |
| Q7P36_000258 | Flavin-containing monooxygenase 3 | 1.5 | Q | no | Cytoplasm |

FC: Fold change. Clusters of Orthologous Groups (COG) categories; C: Energy production and conversion, E: Amino acid transport and metabolism, I: Lipid transport and metabolism, S: Function unknown, and Q Secondary metabolites biosynthesis, transport and catabolism. SP: Signal Peptide. Experiments were performed in biological quadruplicate.

Amongst the 44 significantly up-regulated proteins, four methyltransferases and 19 oxidoreductases were plausible biocatalysts for micropollutant transformation. Of particular relevance, upregulation of members from two oxidoreductase superfamilies was observed: aldo-keto reductases (4 enzymes) and short-chain dehydrogenases/reductases (4 enzymes). In addition, five flavo-enzymes (Baeyer-Villiger monooxygenase Q7P36_007820, 2-hydroxyacid oxidase Q7P36_008506, Sulfide quinone-reductase Q7P36_002133, pyridine nucleotide-disulphide oxidoreductase Q7P36_0010397 and flavin-containing monooxygenase 3 Q7P36_000258) were up-regulated. The remaining enzymes are different types of NAD(P) depend oxidoreductases (Q7P36_004137, Q7P36_006523, Q7P36_004165), and Q7P36_000623 and a single thioredoxin (Q7P36_002807) (Table 1). The majority of the upregulated methyltransferses and oxidoreductases are present in both *C. inversicolor* IBT 42153 (*Cla*5), and *C. fusiforme* IBT 42164 (*Cla*8) (Fig. S6). Notably, the thioredoxin, however, was unique for the *C. allicinum* IBT 42152 (*Cla*4) together with a second less up-regulated counterpart (log_2_= 1.3).

## Discussion

Previous studies have focused on Basidiomycota in the context of micropollutant removal, including compounds in this study(14, 16, 17, 33–35), except for 5-chlorobenzotriazole. However, the present work is the most extensive regarding the span of micropollutants and fungal isolates, e.g. about half of the micropollutants have not previously been tested with ascomycetes. Previous Ascomycota analyses focused on only a few micropollutants, such as diclofenac, which was fully removed in supernatants of a *Phoma* sp. UHH 5-1-03(36) and *Penicillium oxalicum*(37) cultures. The latter strain was also efficient (>80% removal) against ketoprofen(22), in contrast to our strains. In addition, *Trichoderma harzianum* BGP115 and *Trichoderma viride* were shown to remove 71% sulfamethoxazole(38) and 31% ciprofloxacin(39), in contrast to >90% transformation of these antibiotic by *C. allicinum* IBT 42152 (*Cla*4) (Fig. 2). Besides *Trichoderma,* especially *Aspergillus* and *Peniciliium* strains have been shown to remove micropollutants(21, 22, 40). These two, together with eight other Ascomycota genera were included in our initial screening. Strikingly, *Cladosporium* spp., especially, *Cladosporium allicinum* IBT 42152 (*Cla*4), displayed considerable micropollutant transformation (Fig. 1,2).

*Cladosporium* spp. are widespread in marine and terrestrial niches(41) and their general inability to produce mycotoxins makes them well suited to detoxification applications(42–47). In fact, the poorly documented reports on the production of citreoviridin A, viriditoxin, cytochalasin D and citrinin H1 mycotoxins by this genus may be attributed to misidentification or contaminations by mycotoxin-producing species. *Cladosporium* harbors species that can grow at oligotrophic conditions and at fluctuating temperatures(48), which together with their resistant to irradiation(49), supports their robustness in bioremediation applications.

Interestingly, *C. allicinum, C. inversicolor*, and *C. fusiforme* were isolated from soil contaminated with trichloroethylene (TCE, C_2_HCl_3_), which is consistent with high-efficacy transformation of the chlorinated micropollutants, sertraline and 5-chlorobenzotriazole and the previously demonstrated removal of chlorinated pesticides by *C. cladosporioides* and *C. herbarum* (50, 51). *Cladosporium* species were also reported as excellent degraders of polycyclic aromatic hydrocarbons (PAH)(52, 53). However, this is the first report that demonstrates the removals of the six micropollutants 5-methyl-1H-benzotriazole, 5- chlorobenzotriazole, ciprofloxacin, sulfamethoxazole, sertraline, and citalopram by *Cladosporium* members. These micropollutants share at least two aromatic rings, 1-3 nitrogen atoms and are halogenated, except for 5-methyl-1H-benzotriazole and sulfamethoxazole. We identified two transformation products for each of the corrosion inhibitors 5-methyl-1H-benzotriazole and 5- chlorobenzotriazole and a single transformation product for the antidepressant citalopram for all three species, except for *C. allicinum* IBT42152 (*Cla*4) on 5-chlorobenzotriazole, where no products were detected (Fig. 4). The oxygenations (5-methyl-1H-benzotriazole and citalopram) and double methylation and carboxylation (5-chlorobenzotriazole) mediated by *Cladosporium* are different from conjugation to a tetrose sugar and methylation of benzotriazoles, including 5-methyl-1H-benzotriazole, reported for the Basidiomycota model *T. versicolor*(54). Another Basidiomycota, *Pleurotus ostreatus,* was reported to transform 50% of citalopram, resulting in four products corresponding to de-methylation and oxidations, either via oxygenation and/or dehydrogenations(55). The multiple transformation products and modifications, as compared to the fewer products with minimal modifications by the *Cladosporium* spp. is suggestive of different strategies to target micropollutants. In our study, no transformation products were detected for ciprofloxacin, sulfamethoxazole, and sertraline, which may have undergone complete degradation, which is similar to the degradation of the pesticide chlorpyrifos and its hydrolysis product 3,5,6-trichloro-2-pyridinol (TCP) by *Cladosporium cladosporioides* Hu-01(51). Collectively, these findings suggest that *Cladosporium* mediates complete or extensive degradation of certain xenobiotics.

In Basidiomycota, both networks of extracellular oxidoreductases, potentially involved in lignocellulose processing, and cytochrome P450 monooxygenases that catalyse hydroxylation reactions of carbon-carbon bonds have been proposed as key strategies for micropollutants transformation(56). Thus, extracellular high-redox potential laccases were shown to transform distinct micropollutants(57). Laccases from *Cladopsporium* were suggested to transform polychlorinated biphenyls via a redox mediator(58), which was not observed in this study (Table S5), despite the observed laccase-like activities (Fig. 1). Laccase selectivity has been observed, *e.g. Phoma* sp. UHH 5-1-03 cell-free laccase-containing culture supernatants could not remove sulfamethoxazole, as opposed to diclofenac and bisphenol A(36). Moreover, the transformation of the polycyclic compound anthracene by ascomycetes(59) did not correlate to extracellular enzyme activity, including laccase. Collectively, these findings support an intracellular micropollutant transformation machinery for Ascomycota.

Intracellular transformation of PAH(60, 61), triphenyl phosphate(62), and metronidazole(63) by Ascomycota has been porposed. In ascomycetes, xenobiotics are generally metabolized by phase I and phase II detoxification enzymes. Phase I transformations involve oxidations by cytochrome P450 monooxygenases and epoxide hydrolases, which introduce a hydroxy- and epoxy- group in their substrates, respectively(64). Neither P450 nor epoxide enzymes were overexpressed, however, a single P450, (Q7P36_000806), was down-regulated in our work. Other Phase I enzymes include carbonyl-reducing aldo-keto reductases and short-chain dehydrogenases/reductases(65). Four aldo-keto reductases, which were among the top up-regulated proteins in the proteome analysis (Table 1), may play a role in micropollutant transformation (Fig. 7), as demonstrated for a recombinant enzyme on a cancer drug(66). Short-chain dehydrogenases/reductases constitute a large family of NAD(P)(H)-dependent oxidoreductases(67), members of which were upregulated in *Aspergillus* in the presence of PAH(60). Four up-regulated members of this family (Table 1) hint the involvement in micropollutant transformation in *Cladosporium*. Similarly, the up-regulation of a putative flavo-monooxygenase 3 (FMO3) was observed (Q7P36_000258). The human FMO3 is non-CYP P450 liver enzyme catalysing the *N*-oxidation of trimethylamine to trimethylamine *N*-oxide using NADPH, and the monooxygenation of nitrogen- or sulphur-containing xenobiotics(68). Thus, this enzyme is a plausible candidate to perform the oxygenation of the products identified by the network analysis (Fig. 4).

**FIG 7:**
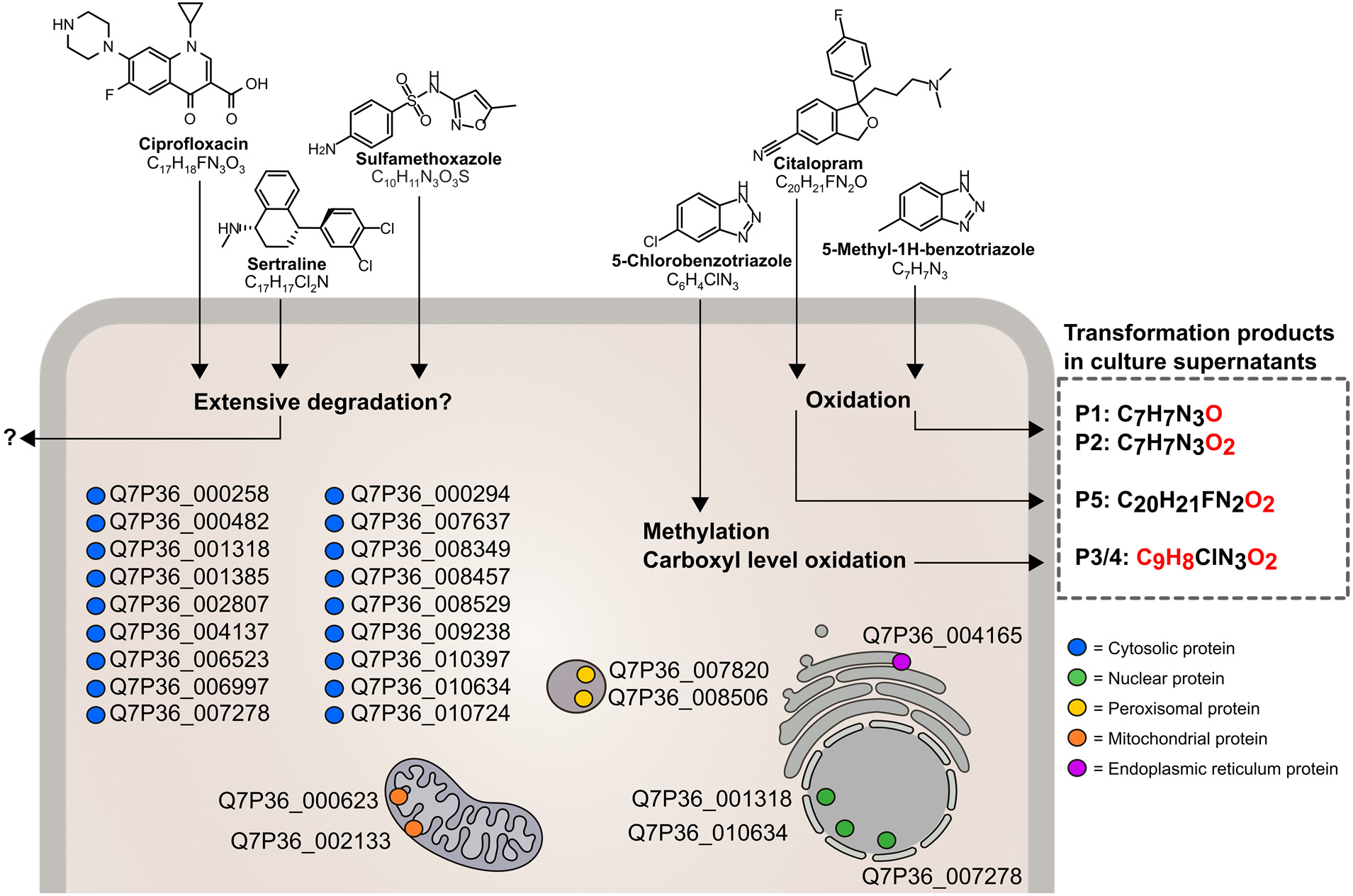
Proposed model for the metabolism of micropollutants by *Cladosporium* species. The six micropollutants 5-methyl-1H- benzotriazole, 5-chlorobenzotriazole, ciprofloxacin, sulfamethoxazole, sertraline, and citalopram are likely internalized into the fungal cell. 5-methyl-1H-benzotriazole and citalopram are oxidized, while 5-chlorobenzotriazole is double methylated and oxidized to the carboxyl level. Certain transformation products are likely effluxed from the cell via unknown systems, based on their detection in the culture supernatant. The transformation mechanism for ciprofloxacin, sulfamethoxazole, and sertraline could not be assigned, but the lack of transformation products indicates more extensive degradation. The oxidoreductases and methyltransferases upregulated log_2_ ≥ 1.5 in the proteomic are indicated in the model, due to their potential involvement in the conversions of the investigated micropollutants.

Methyltransferases (phase II) play pivotal roles in the methylation of xenobiotics by transferring a methyl from a donor to substrates(69), and these enzymes have been shown to be up-regulated(62) and important for biotransformation of xenobiotics by distinct Ascomycota(70). Our proteomic study showed that 4 methyltransferases were among the top up-regulated proteins (Table 1) in the presence of the micropollutants, consistent with double methylation of 5-chlorobenzotriazole (Fig. 4). These finding indicate the involvement these methyltransferases in the modification of distinct micropollutants, along their transformation itinerary (Fig. 7).

Key concerns regarding persistent micropollutants, is their potential cytotoxicity. Indeed, we showed the inhibition of the *Cladosporium* strains by 5-methyl-1H-benzotriazole, 5-chlorobenzotriazole, sulfamethoxazole, and sertraline (Fig. 3B,C). Stress from exogenous micropollutants can lead to overproduction of reactive oxygen species (ROS)(71), which mediate oxidative damage and the aberrant disulphide bridge formation in proteins. Thioredoxin is a ubiquitous redox regulator, which protects cytosolic proteins from oxidative stress(72, 73). Two thioredoxins were up-regulated (log_2_>1.3), suggesting that the thioredoxin in *C. allicinum* IBT 42152 (*Cla*4) protects against oxidative stress. Similar observations were observed for *Aspergillus sydowii* FJH-1(62). The two other thioredoxins, which were lacking in the two other Cladosporium strains (Fig. S7), may also contribute to the resilience of *C. allicinum* IBT 42152 (*Cla*4) against oxidative stress. Efficient systems to alleviate oxidative damage, merit additional attention in fungal micropollutant transformation efficacy. Finally, transport systems that mediate internalization of micropollutants, and their potential externalisation of modified/detoxified forms (Fig. 3D), remain an uncharted territory. Future studies are needed to focus on the transport aspects.

### Conclusions

Our study discovered potent micropollutant transformation and detoxification capabilities for three previously not explored *Cladosporium* species. We also provided the first genomes for all these species, as a useful resource for the community for future comparative genomics and molecular work on these underexplored taxonomic groups within bioremediation. Proteomics and metabolomics data supported the involvement of specific intracellular oxidoreductases and methyltransferases in micropollutant biotransformation. However, knock-out or recombinant expression of these enzyme is needed to confirm their role in transformation of the micropollutants. Our study provides novel insight into the potential of Ascomycota for micropollutant transformation and sets the stage for further investigations to evaluate *Cladosporium* species and their enzymes in bioremediation of wastewater.

## Materials and methods

### Chemicals

The micropollutants (diclofenac sodium salt, bicalutamide, benzotriazole, 5-chlorobenzotriazole, 5- methyl-1H-benzotriazole, clofibric acid, sertraline hydrochloride, iohexol, ketoprofen, carbamazepine, clarithromycin, venlafaxine hydrochloride, erythromycin, atenolol, metoprolol tartrate salt, hydrochlorothiazide, bezafibrate, mefenamic acid, ciprofloxacin, azithromycin, citalopram hydrobromide, and sulfamethoxazol) were purchased from Merck (Darmstadt, Germany) (Table S1). Acetonitrile, formic acid and methanol used for micropollutant analysis were HPLC-gradient grade (Merck, Germany).

### Strains

The 53 fungal strains investigated in the present study were from the strain collection of the Department of Biotechnology and Biomedicine (previously Institute for Biotechnology, IBT), Technical University of Denmark. The selected strains were isolated from trichloroethylene (TCE) polluted soil (Vedbæk, Denmark) or from wastewater at the Lynetten wastewater treatment plant (Copenhagen, Denmark) (Table S2). Conidia from the fungal strains IBT 31311-31408 were propagated on Czapek yeast extract agar (CYA) and conidia from the fungal strains IBT 42148-42169 were propagated on malt extract agar MEA, incubated at 25 °C for 7 days. Fresh spores for the inoculum were kept in 0.9% (w/v) NaCl, 0.0005% (v/v) Tween 80 at 5 °C.

### Growth

Initial screening of fungal strains for removal of micropollutants was performed in 10 mL glass tubes containing 6 mL liquid media; Czapek yeast extract medium (per liter: 5 g yeast extract, 35 g Czapek dox broth, 1 mL trace element solution), malt extract medium (per liter: 20 g malt extract broth, 1 mL trace element solution), or minimal medium (MM) (per liter 1 g KH_2_PO_4_, 1 g KNO_3_, 0.5 g MgSO_4_, 0.5 g KCl, 0.2 g glucose, 0.2 g sucrose). The trace element solution consisted of 1 g L^-1^ ZnSO_4_·7H_2_O, 0.5 g L^-1^ CuSO_4_·5H_2_O. The selected 53 strains were designated with the first three letter of their genus affiliation (based on phenotypic typing) and a number and the list of these strains and their corresponding IBT collection numbering is shown in Table S2. *Aspergillus (Asp)* and *Penicillium (Pen)* strains were grown in Czapek yeast extract medium and *Cladosporium (Cla), Fusarium (Fus), Stachybotrys (Sta), Geotrichum (Geo), Phoma (Pho), Monilochaetes (Mon), Onychocola (Ony),* and *Epicoccum (Epi)* strains were grown in malt extract medium. The 22 micropollutant blend (Table S1) was made from the stocks solutions of each micropollutant (5 mg mL^-1^ dissolved in MeOH) and added to the media, resulting in a final concentration of 1 mg L^-1^ of each pollutant. Cultures were inoculated with 300 µL spore suspension in three biological replicates and incubated in the dark for 12 days in a Kühner ISF-1-W orbital shaker (20 °C, 75 rpm).

A time course experiment was performed in larger scale (250 mL baffled Erlenmeyer flasks with 50 mL MM) containing a mixture of all micropollutants at a final concentration of 1 mg L^-1^ of each. The flasks were inoculated with 1·10^6^–10·10^6^ conidia in three biological replicates and incubated in the dark for 15 days under agitation in a Kühner ISF-1-W orbital shaker (20 °C, 100 rpm). Abiotic controls comprised medium without inoculation, but otherwise under the same experimental conditions. To evaluate the removal of micropollutants from culture supernatants, 2 mL culture suspensions were centrifuged (10 min, 20,000 *g*) to remove fungal biomass and 1.5 mL supernatant was transferred into a glass tube with 0.4 mL acetonitrile (ACN) to quench bioactivities. All samples were stored at −20 °C prior to analysis by High-Performance Liquid Chromatography-Electrospray Ionization Tandem Mass Spectrometry (HPLC-ESI MS/MS) for micropollutant quantitation.

### Extracellular laccase activity

Laccase-like activity was assayed by monitoring the oxidation of 0.5 mM ABTS (2,2’azino-bis(3- ethylbenzothiazoline-6-sulphonic acid) in assay buffer (0.1 M sodium acetate buffer, 0.001% (v/v) triton X-100, pH 4.5) at 420 nm (*ε*_420_ = 36,000 M^-1^ cm^-1^), at 25 °C for 1-15 hours. The assay was performed in a microtiter plate with 40 µL supernatant, 245 µL buffer and 15 µL 10 mM ABTS. One unit (U) was defined as the amount of enzyme activity required to oxidize 1 µmol ABTS per min.

### Evaluation of the ability of fungal secretomes in culture supernatants to transform micropollutants

Fungal growth was performed as described above in 250 mL baffled Erlenmeyer flasks, but in 90 mL MM instead of 50 mL. Supernatants (35 mL) were removed at day 6 and day 10 and filtered through a Miracloth (Merck Milipore, Carrigtwohill, Ireland) into a clean tube, centrifuged (15 min, 4000 *g*) and re-filtered to remove fungal spores, using a 0.45 µm Q-Max® Syringe filter (Knebel, Denmark). Aliquots (0.5 mL) were collected and the rest of the supernatants were concentrated from 35-fold to 1 mL using an Amicon®Ultra Centrifugal Filters, cut off 10 kDa (Merck Milipore). Half of the filtered supernatant and half of the concentrated supernatant were heat treated (15 min, 90 °C) to inactivate potentially secreted enzymes to serve as negative controls. The transformation assay was carried out in 1.4 mL, 10 mg mL^-1^ of each micropollutant, 50 mM acetate buffer, pH 5 and started by adding 100 µL of either the x1- or the x35-fold concentrated culture supernatant. Samples were incubated at 37 °C for 3 days, stopped by addition of 400 µL ACN and stored at −20 °C until further analysis. The experiment was performed in biological triplicates.

### Analysis of micropollutant transformation

The quantitative analysis of the micropollutants after fungal growth was carried out by HPLC (Agilent, 1290 Infinity, USA) coupled with triple quadrupole mass spectrometer (Agilent, 6470 series, USA).

Prior to analysis the supernatants were diluted 40 times with ultrapure water in a glass tube (212−0016, DURAN®, VWR, Denmark. For the analysis, 900 µL of each diluted sample and 100 µL of the internal standard solution (500 µg L^-1^: 5-Methyl-1h-benzotriazole-d6, atenolol-d7, atrazine-d5, azithromycin-d3, bezafibrate-d4, ciprofloxacin-d8, carbamazepine-d8, citalopram-d6, diclofenac-d4, iohexol-d5, ketoprofen-d3, metoprolol-d7, sertraline-d3, sulfamethoxazole-C6 and venlafaxine-d6)were transferred into an HPLC vial (8010-0543, Agilent, Germany). 10 μL of mixture from the HPLC vial was injected into HPLC (Agilent 1290 Infinity, USA), coupled with a triple-quadrupole mass spectrometer (Agilent 6470 series, USA). The chromatographic separation was performed using a C18 column (2.1x50 mm, 1.8 µm, Eclipse Agilent, USA), HPLC column oven temperature = 35 °C, and a constant flow rate = 0.5 mL min^-1^ in ESI+ mode or 0.6 mL min^-1^ in ESI- mode. A gradient of 0.1% (v/v) formic acid (FA) in ultrapure water (Eluent A) and 0.1% (v/v) FA in ACN (Eluent B) was used in ESI+ mode, while a 10 mM ammonium acetate in ultrapure water (Eluent A) and 10 mM NH_4_OAc in 90% ACN/10% (both v/v)ultrapure water (Eluent B) was used in ESI- mode.

Dynamic multiple reaction monitoring (MRM), which is able to automatically separate the transitions of contaminants into multiple MRM tables according to the retention time window for each transition, was implemented for ESI+ modes. For the analysis under ESI+ mode, the HPLC gradient was initiated with 0% B for 2 min, followed by increasing gradient B to 15%, 20%, 36%, 45%, 60%, and 75% after 2 min, 3 min, 2 mins, 1 min, 2 min and 2 min, respectively. Before each sample injection of 10 µL, initial gradient conditions were re-established for 3 min. Nitrogen was used as the sheath gas as well as the collision gas, while for the analysis under ESI- mode, a normal MRM method was implemented. The HPLC gradient was initiated with 0% B, followed by a linear increase of gradient B up to 60% within 8 min. Initial gradient conditions were also re-established for 3 min. For both ESI+ and ESI-, sheath gas flow=12 L min^-1^, T=400 °C, gas flow=10 L min^-1^ with the temperature of 260 °C. The voltage for the capillary was 4500 V, while the voltage for ESI+ and ESI- was 500 and 0 V, respectively.

The micropollutant removal percentage was calculated using:

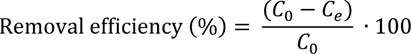

where, *C*_0_ is the initial concentration of pollutants in the culture supernatant (mg L^-1^) and *C_e_* is the final concentration of the pollutants (mg L^-1^).

### Biosorption

Sorption of the micropollutants onto fungal mycelia was analysed to estimate its contribution to the observed removal of the micropollutants from culture supernatants. After 15 days culturing with micropollutants in MM, fungal biomass was separated from the supernatant by filtration using a miracloth, washed with 20 mL 0.9% (w/v) NaCl and transferred to a 50 mL Falcon tube. Subsequently, 3 mL ACN were added and the tubes were vortexed for 30 s, sonicated for 10 min, and agitated for (1 hour, 300 rpm) using an Eppendorf Thermoshaker at room temperature. Subsequently, water was added to reconstitute to the volume of the culture suspension before filtration. Next, 2 mL culture suspensions were centrifuged (10 min, 20,000 *g*), 1.5 mL supernatant was transferred to 0.4 mL ACN and analysed by HPLC-ESI MS/MS. The concentrations of micropollutant identified in the extractions were assumed to be due to sorption onto the fungal biomass.

### Micropollutant toxicity and detoxification

The toxicity of the micropollutants ciprofloxacin, sulfamethoxazole, citalopram, setraline, 5- chlorobenzotriazole, and 5-methyl-1H-benzotriazole was assayed based on growth inhibition by measuring the diameter of the fungal colonies, which were grown on MM agar plates, supplemented either with individual micropollutants or the mixture thereof. Three micropollutant concentrations were applied: 0, 1, and 10 mg L^-1^. The plates were incubated for 11 days at 25 °C and colony diameter (mm) was measured every third day in four biological replicates (n=4).

For evaluating changes in toxicity, the *Cladosporium Cla*4, *Cla*5, and *Cla*8 were grown in MM with a mixture of the micropollutant including ciprofloxacin, sulfamethoxazole, citalopram, setraline, 5- chlorobenzotriazole, and 5-methyl-1H-benzotriazole (1 mg L^-1^ each) and the biomass separated from supernatant as described above. Subsequently, the supernatants were freeze dried and reconstituted in LB medium to a concentration of 2 mg L^-1^ and 20 mg L^-1^ of each micropollutant. To assay if the supernatants from the culture supernatants were detoxified, an overnight culture of a *Escherichia coli* DH5α grown in LB medium was used to inoculate (1% v/v) 200 µL of the LB media, containing the micropollutants as described above, and grown for 12 hours at 37 °C. The experiment was performed in biological triplicates (n=3), which were analyzed in three technical triplicates for each biological replicate.

### Identification of transformation products in culture supernatants

The micropollutant transforming strains *Cla*4, *Cla*5, and *Cla*8 were individually grown in MM, supplemented with a mixture of the following pollutants: ciprofloxacin, sulfamethoxazole, citalopram, setraline, 5-chlorobenzotriazole, and 5-methyl-1H-benzotriazole as described above. Two controls were included - an abiotic control (no inoculum), and one without the micropollutant mixture. The experiments were performed in four independent biological replicates (n=4). After 15 days, 2 mL culture suspensions were centrifuged (10 min, 20,000 *g*) to remove fungal biomass and 1.5 mL supernatant was transferred to 0.4 mL ACN. Samples were subsequently concentrated 12-fold using an Eppendorf Concentrator Plus (45 °C). Finally, ACN was added to (20% v/v) prior to the LC-ESI MS/MS analysis.

The samples were analysed using an Agilent 1290 Infinity II UHPLC (Agilent Technologies; Santa Clara, CA) equipped with an Agilent Poroshell 120 Phenyl Hexyl column (1.9 µm, 150 × 2.1 mm) coupled to an Agilent 6545 QTOF. Elution was performed using a gradient of 20 mM FA in LC-MS grade water (Eluent A) and 20 mM FA in LC-MS grade ACN (eluent B). The gradient started at 10% eluent B, increasing to 100% over 10 min, held at 100% for 2 min at a flow rate= 0.35 mL min^-1^ and a column temperature of 60 °C. Ionization was achieved using ESI and mass spectra were obtained in the m/z =100-1600 range, acquired at a rate of 10 scans s^-1^. All MS analyses was undertaken with internal standards (hexakis-(1H, 1H, 3H- tetrafluoropropoxy)-phosphazine, HP-0921) for spectra calibration.

An untargeted approach using feature based molecular networking was used to identify transformation products. The acquired mass-spectra were imported into MZMine 3(74) for feature detection and alignment. The results were exported to the GNPS web platform(75) for feature based molecular network analysis and the GNPS-generated results were visualized and analyzed in Cytoscape(76) for identification of transformation products. Features that formed clusters with the spiked pollutants, as well as having an increase in abundance over the incubation days were considered features of interest and potential transformation products.

### Genome sequencing

The tree *Cladosporium* strains *Cla*4, *Cla*5, and *Cla*8 were cultivated on Potato Dextrose Agar plate. Three 10 µL loops of the mycelia (about 30 µg) were transferred to 200 mL modified and diluted LB medium (1 g tryptone, 1 g NaCl and 1 g yeast extract). The flasks were grown for 14 days, at 20 °C, 80 rpm in the dark. Mycelia were separated using a Miracloth, washed twice with water followed by lyophilization using a freeze-dryer and grinding in a mortar. Genomic DNA was extracted from the mycelial powder using the phenol-chloroform method(77). Purification of the DNA and removal of small fragments were performed according to same protocol. The quality and quantity of the extracted DNA were evaluated using

NanoDrop One (ThermoFisher), Qubit 3.0 (Invitrogen) with Qubit dsDNA HS Assay Kit, and 2200 TapeStation (Agilent) with Genomic DNA ScreenTape Analysis according to the manufacturer’s instructions. A library was constructed using the Native barcoding genomic DNA (EXP-NBD104, EXPNBD114, and SQK-LSK109) from Oxford Nanopore Technologies (Oxford, UK) and sequenced on a R9.4.1 flowcell.

### De Novo assembly, genome annotation and comparative genomics

The raw data were basecalled, demultiplexed, and adapters were removed using Guppy version 6.1 (https://github.com/nanoporetech/pyguppyclient) in GPU mode using thedna_r9.4.1_450bps_hac.cfg model. The reads were filtered using Filtlong version 0.2.0 (https://github.com/rrwick/Filtlong) to a minimum length of 10 kb and a minimum basecall quality of 80 (Q7). Minimap2 version 2.17(78) and Miniasm version 0.3(79) were used to create the assembly, which subsequently were polished using Racon version 1.3.3(80) with default settings and two rounds of Medaka version 1.6 (https://github.com/nanoporetech/medaka) with default settings. The completeness was assessed with Benchmarking Universal Single-Copy Orthologues (BUSCO) version 5(81) using the Ascomycota version 10 BUSCO data set. The genome was annotated using AUGUSTUS version 3.4.0(82) with *Aspergillus nidulans* as model organism, using EggNOC-mapper version 2.1.9(83) and InterPro version 90.0(84) for functional annotation. Noncoding RNA genes were predicted using Barrnap version 0.9 (https://github.com/tseemann/barrnap). The internal transcribed spacer regions (ITS)(85), partial actin (*act*)(86), and translation elongation factor 1-alpha (*tef1*)(86) genes in the three assembled genomes were used for species assignment by comparison to fungal orthologues using the BLASTn tool of the NCBI GenBank nucleotide database. The assembled and annotated genomes were deposited at NCBI under BioProjectPRJNA980560. For comparative analyses, the 23 available *Cladosporium* genomes on NCBI were downloaded and annotated using AUGUSTUS as described above. *Aspergillus nidulans* FGSC A4 proteome was included as a reference. CAZymes were annotated for all genomes with dbCAN2(87). FastANI(88) was used for whole genome Average Nucleotide Identity (ANI) analysis.

### Proteomics sample preparation and LC-MS

For the proteomic analyses, the *Cla*4 strain was grown in 50 mL MM for 7 days in four independent biological replicates (n=4) in a medium supplemented with a mixture of the micropollutants ciprofloxacin, sulfamethoxazole, citalopram, setraline, 5-chlorobenzotriazole, and 5-methyl-1H-benzotriazole (1 mg mL^-1^ each) and a control culture lacking the micropollutants was included. The biomass was separated from the supernatant using a Miracloth. For secretome analyses, supernatants were filtered (0.22 μm PES syringe filters) and proteins from 10 mL culture filtrates were precipitated by addition of acetone to a final concentration of 80% (v/v) and incubation over night at −20 °C. Next, the proteins were pelleted by centrifugation (10 min, 18,000 *g*, 5 °C) and the supernatants discharged. The pellets were air-dried for 30 min, dissolved in 60 µL lysis buffer (6 M Guanidinium Hydrochloride, 10 mM Tris(2- carboxyethyl)phosphine hydrochloride, 40 mM 2-Chloroacetamide, 50 mM HEPES pH 8.5), boiled (5 min, 95 °C) and sonicated (5x60 s, 4 °C) (Bioruptor, Diagenode). For the intracellular proteomes, the mycelial biomass was flash frozen in liquid nitrogen, lyophilized and subsequently ground in a mortar into a fine powder. The powder (2 mg) was dissolved in 75 µL lysis buffer, boiled (5 min, 95 °C) and sonicated (5x60 s, 4 °C) (Bioruptor, Diagenode). Lysates from the secretomes and intracellular fractions were centrifuged (10 min, 14,000 *g*, 4 °C) and soluble protein concentrations were determined (Bradford assay, Thermo Fisher Scientific). For digestion, 10-20 µg protein were diluted 1:3 with 50 mM HEPES, 10% (v/v) ACN, pH 8.5 and incubated with LysC (MS grade, Wako) in a ratio of 1:50 (LysC:protein) for 4 h at 37 °C. Subsequently, samples were diluted to 1:10 with 50 mM HEPES, 10% (v/v) ACN, pH 8.5 and further digested with trypsin (MS grade, Promega) in a ratio of 1:100 for 20 h at 37°C. The digestion was stopped by the addition of 2% (v/v) trifluoroacetic acid (TFA) in a ratio of 1:1. Samples were desalted using two-three discs of C18 resin packed into a 200 µL tip and activated by successive loading of 30 µL MeOH, 30 µL 80% (v/v) ACN, 0.1% (v/v) FA and 2x 30 µL of 3% (v/v) ACN, 1% (v/v) FA. The samples were loaded in steps of 50 µL. After loading, tips were washed three times with 100 µL 0.1% (v/v) TFA and peptides were eluted in two steps with 30 µL each of 40% ACN (v/v), 0.1% (v/v) FA into a 0.5 mL Eppendorf LoBind tube. Eluted peptides were dried in an Eppendorf Speedvac (1 h, 60 °C) and reconstituted in 12 µL 2% (v/v) ACN, 1% (v/v) TFA with iRT peptides prior to mass spectrometry analysis. Peptides (500 ng) were loaded on the mass spectrometer by reverse phase chromatography through an inline 15 cm C18 column (Thermo EasySpray ES804) connected to a 2 cm long C18 trap column (Thermo Fisher 164946) using a Thermo Easy-nLC 1200 HPLC system. Peptides were eluted at 250 nL min^−1^ over 140 min with a five-step ACN gradient: 0–85 min gradient of 10–23%, 85–115 min 23–38%, 115–125 min 38–60%, 125–130 min 60–95%, and 130–140 min 95%. Analysis was performed on a Q-Exactive Orbitrap (Thermo Fisher Scientific) run in a data-dependent MS/MS (DD-MS2) topN method, with a loop count of 10. Full MS spectra were collected at 70,000 resolution, with an automatic gain control (AGC) target of 3×10^6^ ions or maximum injection time of 20 ms. Peptides were fragmented via higher-energy collision dissociation (normalized collision energy = 25). The intensity threshold was set to 1.7×10^4^, dynamic exclusion to 60 s and ions with a charge state <2 or unknown species were excluded. MS/MS spectra were acquired at a resolution of 17,500, with an AGC target value of 1×10^6^ ions or a maximum injection time of 60 ms. The scan range was limited from 300–1750 m/z.

### Label-free quantitative proteomics analysis

The proteomics data have been deposited to the MassIVE with the dataset identifier MSV000092869. The raw files were analysed using Proteome Discoverer 2.4 and 3.0 (Thermo Fisher Scientific). Label-free quantitation (LFQ) was enabled in the processing and consensus steps, and spectra were matched against the proteome of *Cla*4. Dynamic modifications were set as oxidation of Met residues, and Acetyl on protein N-termini. Cysteine carbamidomethyl was set as a static modification. The results were filtered to a 1% false discovery rate (FDR). Only proteins for which two unique peptides detected (as defined in Proteome Discoverer) were analysed. Log_2_ fold changes were calculated by comparing the LFQ intensity of each protein in the samples with pollutants to the control samples. Proteins with log_2_ fold changes ≥1.5 or ≤ −1.5 and p-value ≤ 0.05 were identified as up-regulated and down-regulated, respectively. Signal peptides were predicted using SignalP 6.0(89), transmembrane domains were predicted using DeepTMHMM(90), subcellular localization was predicted using DeepLoc 2.0(91) and InterPro 92.0(84) was used to classify proteins into families. The proteins were blasted against UniProtKB/Swiss-Prot and the non-redundant protein sequence database using BLASTp(92).

## Acknowledgements

This work was supported by an Interdisciplinary Synergy Grant from The Novo Nordic Foundation under the grant number NNF18OC0034918.

We thank Associate Professor Jakob Blæsbjerg Hoof (Technical University of Denmark) for discussion on species assignment. We also thank the IBT culture collection and technical assistant Lisette Knoth-Nielsen (DTU-Bioengineering) for the fungal strains and technical support. We are also very grateful for the help of Marie Vestergaard Lukassen and Maike Wennekers Nielsen from the proteomic core facility at the Department of Biotechnology and Bioengineering at DTU for their kind help with the acquisition and analyses of the Proteomics data.

## Author contribution

MLL: Conceptualisation, data curation, formal analysis, investigation, visualisation, writing original draft and review and editing; KT: Data curation, formal analysis, methodology, writing-review & editing; TS: data curation, formal analysis, writing - review & editing; AJA: formal analysis, methodology, writing - review & editing; CHN: Conceptualisation, funding acquisition, project administration, writing - review & editing; BA: Conceptualisation, methodology, resources, writing - review & editing; JF: conceptualisation, methodology, resources, writing - review & editing; TES: Methodology, resources, writing - review & editing; HRA: Conceptualization, methodology, resources, writing - review & editing; MAH: Conceptualization, funding acquisition, project administration, resources, supervision, writing original draft and review & editing

## Data availability

Data have been deposited as follows: The assembled and annotated genomes of the three *Cladosporium* species were deposited at NCBI under BioProjectPRJNA980560. The proteomics data have been deposited to the MassIVE with the dataset identifier MSV000092869

## Supplemental Material Figures

**FIG S1:**
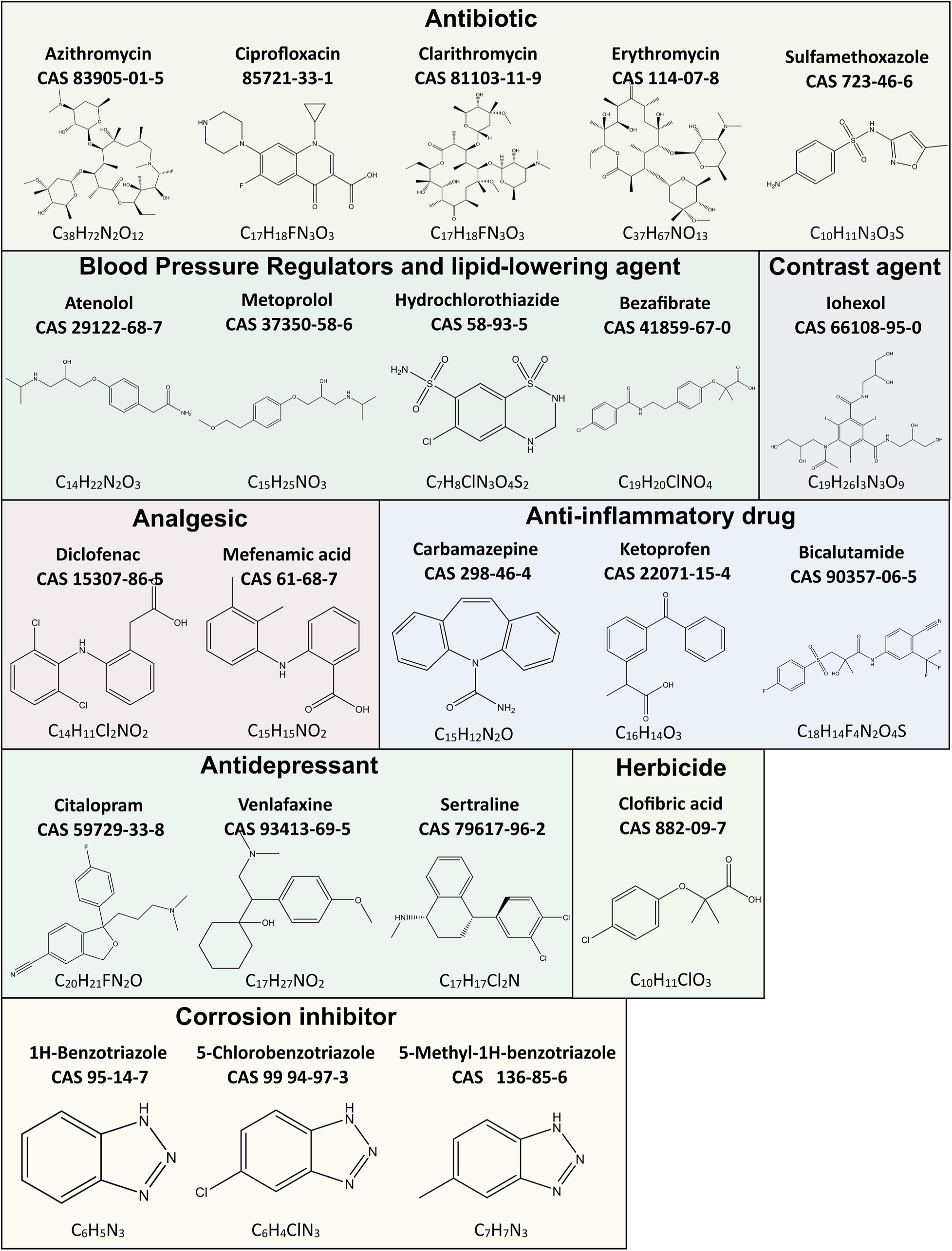
Overview of the micropollutant used in the study

**FIG S2:**
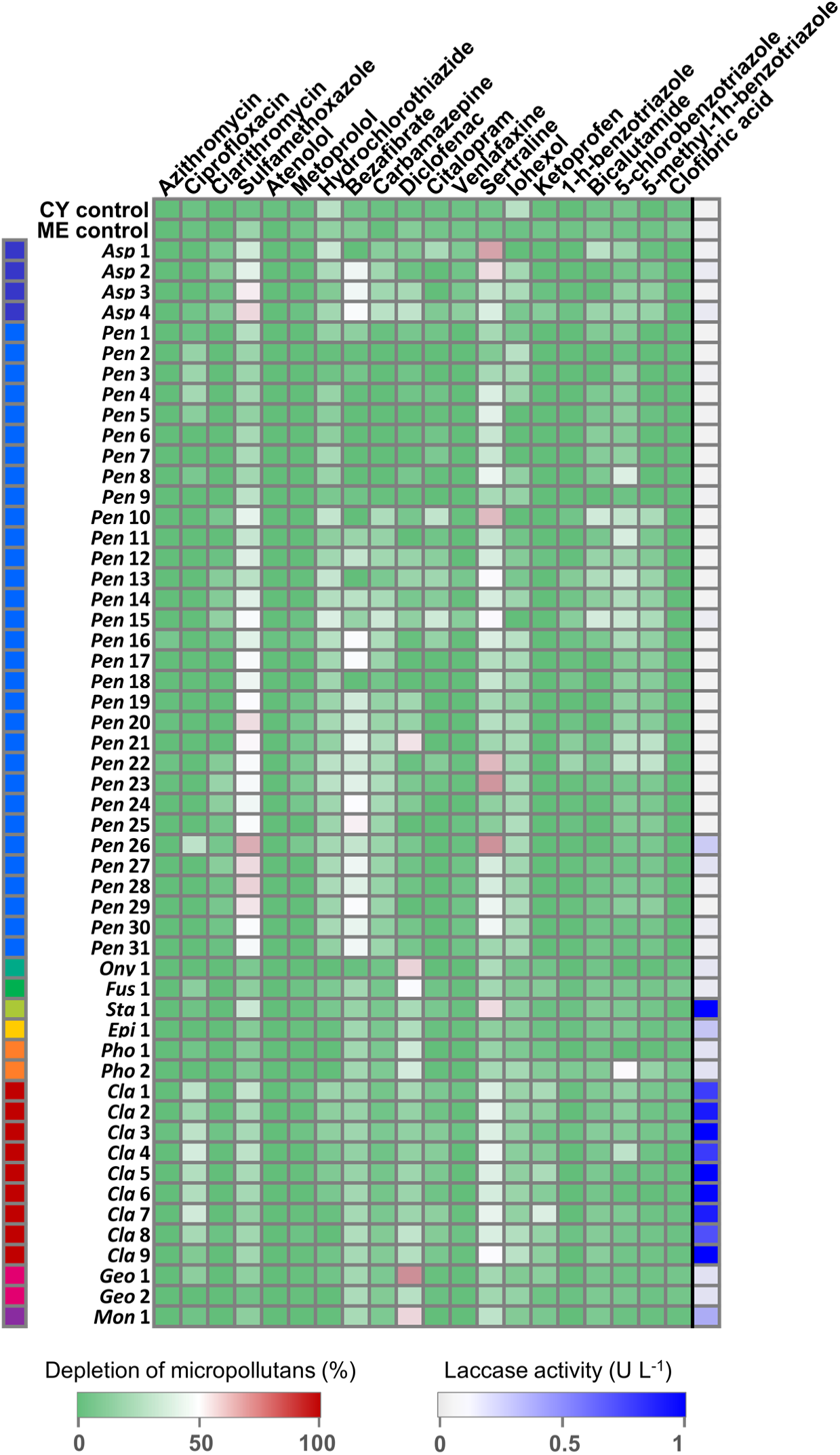
A Heat map of micropollutants removal and laccase-like activities by the Ascomycota isolate panel measured after 12 days incubation in nutrient rich media. The results are shown in percentage relative to an abiotic control. The experiment was performed with three biological replicates.

**FIG S3:**
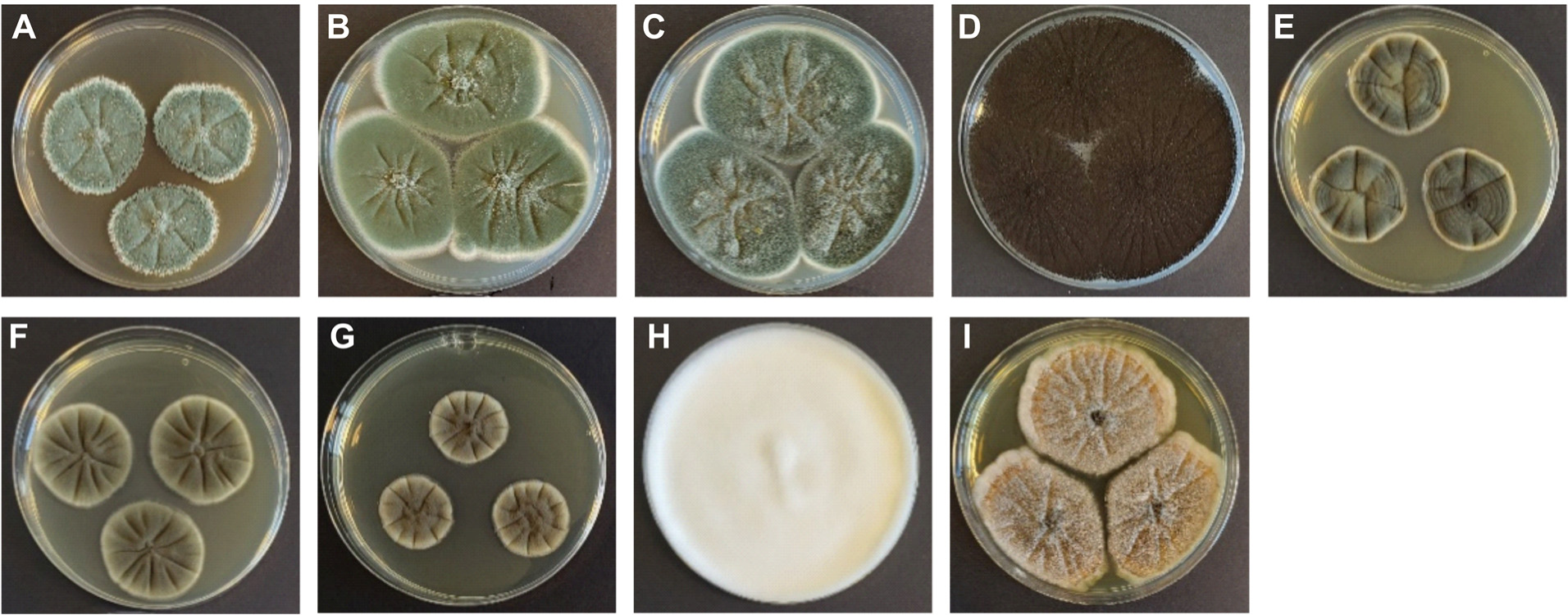
Fungal strains that were shown to be efficient in removal of micropollutants. A) *Pen*4, B) *Pen*9, C) *Pen*22, D) *Asp*4, E) *Cla*4, F) *Cla5,* G) *Cla8,* H) *Geo*1, I) *Mon*1. A-C were grown on Czapek yeast agar (CYA) plates and D-I were grown on malt extract agar (MEA) plates.

**FIG S4:**
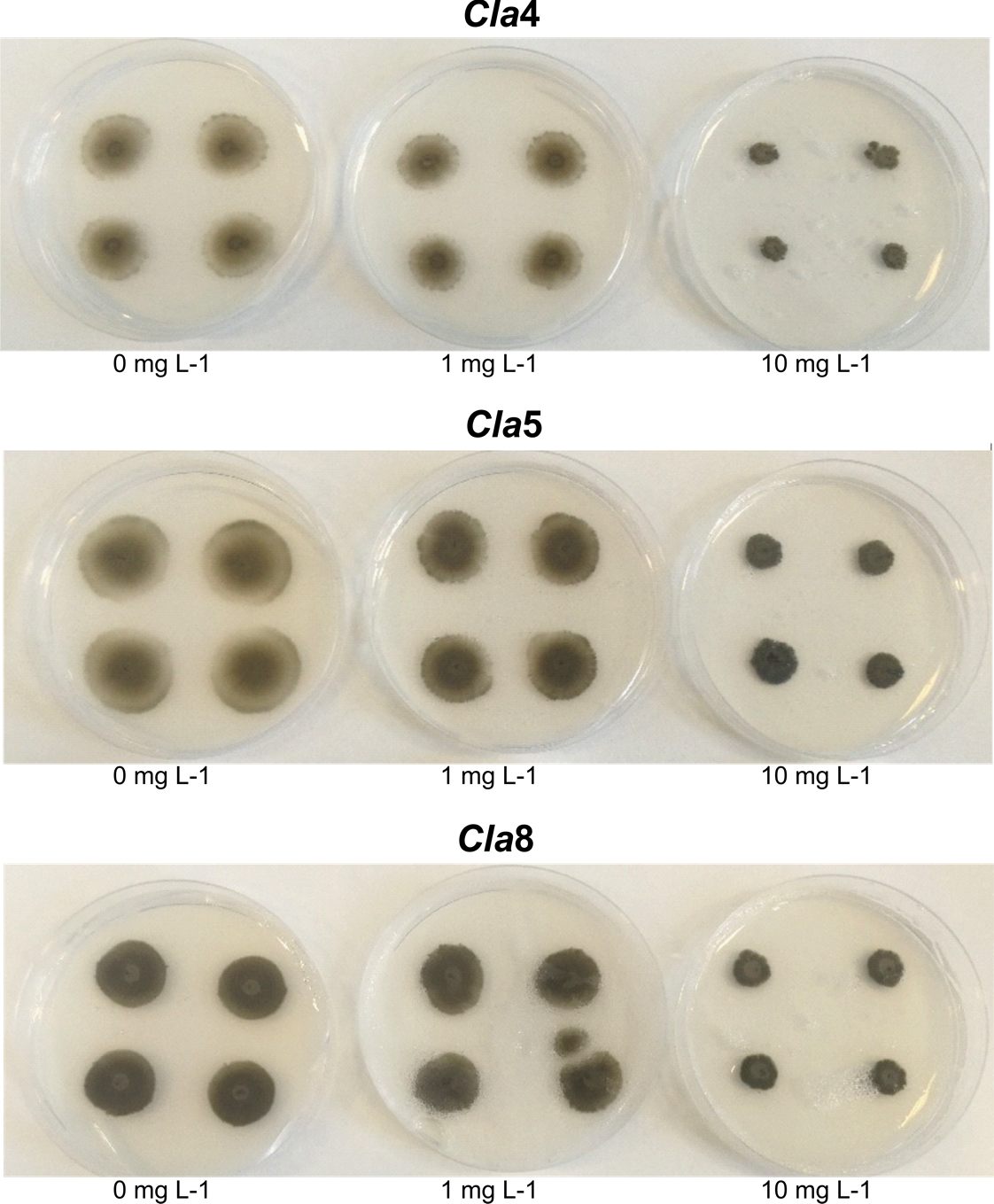
Inhibition of fungal growth by a mixture of the micropollutants. Three *Cladosporium*; *Cla*4*, Cla*5*, and Cla*8 were grown on MM agar plates with a mixture of the micropollutants including 5-methyl-1H-benzotriazole, 5-chlorobenzotriazole, ciprofloxacin, sulfamethoxazole, sertraline, and citalopram. The concentration of each micropollutant in the mixtures was 0 mg L^-1^, 1 mg L^-1^, or 10 mg L^-1^.

**FIG S5.**
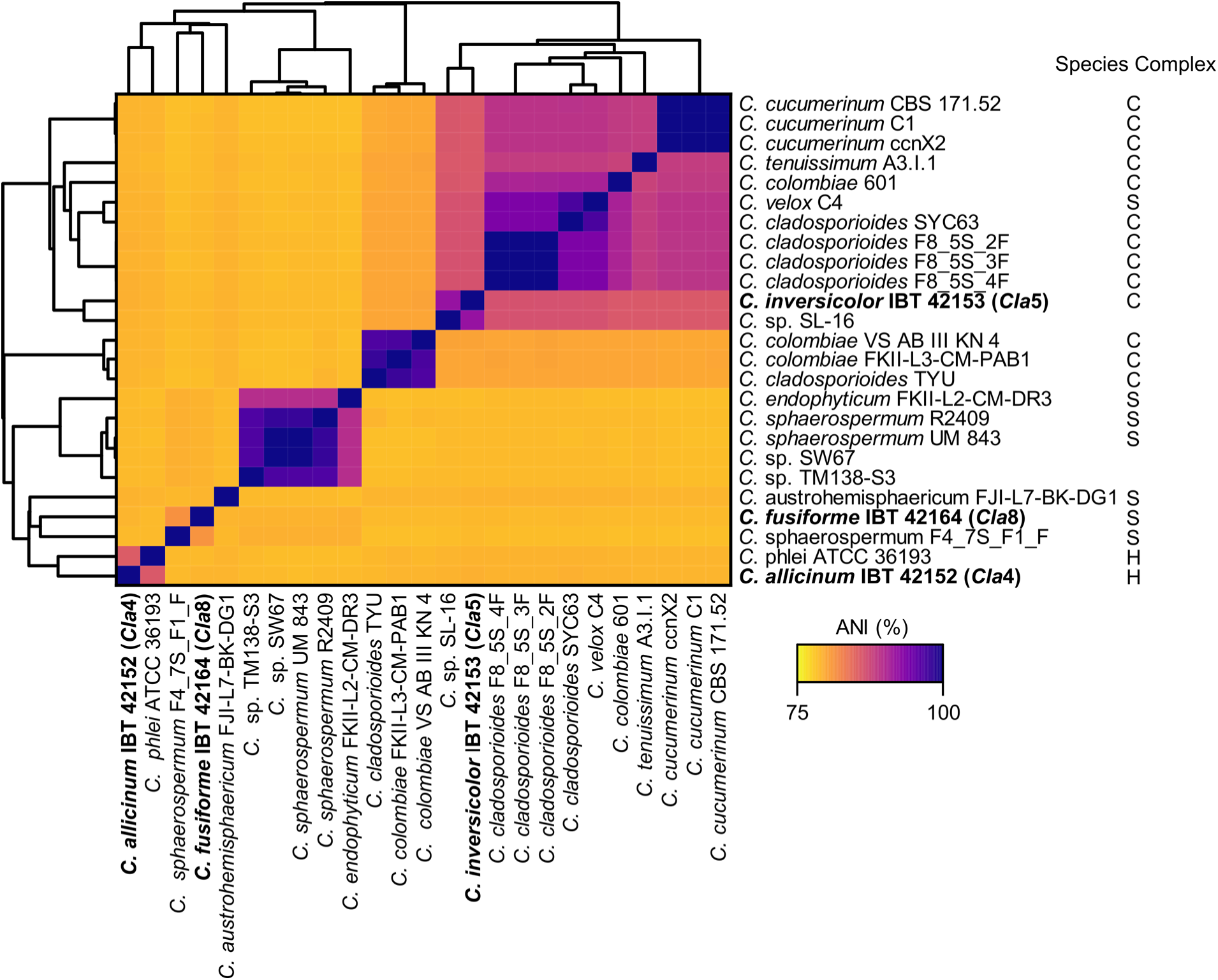
Heatmap of genomic average nucleotide identity (ANI) of pairwise whole genome comparison of *Cladosporium* species. The depicted heatmap is a matrix generated by FastANI and clustering was based on a Euclidean distance. The species complexes of the individual strains are shown to the right, C: *C. cladosporioides*, S: *C. sphaerospermum*, H: *C. herbarum*.

**FIG S6.**
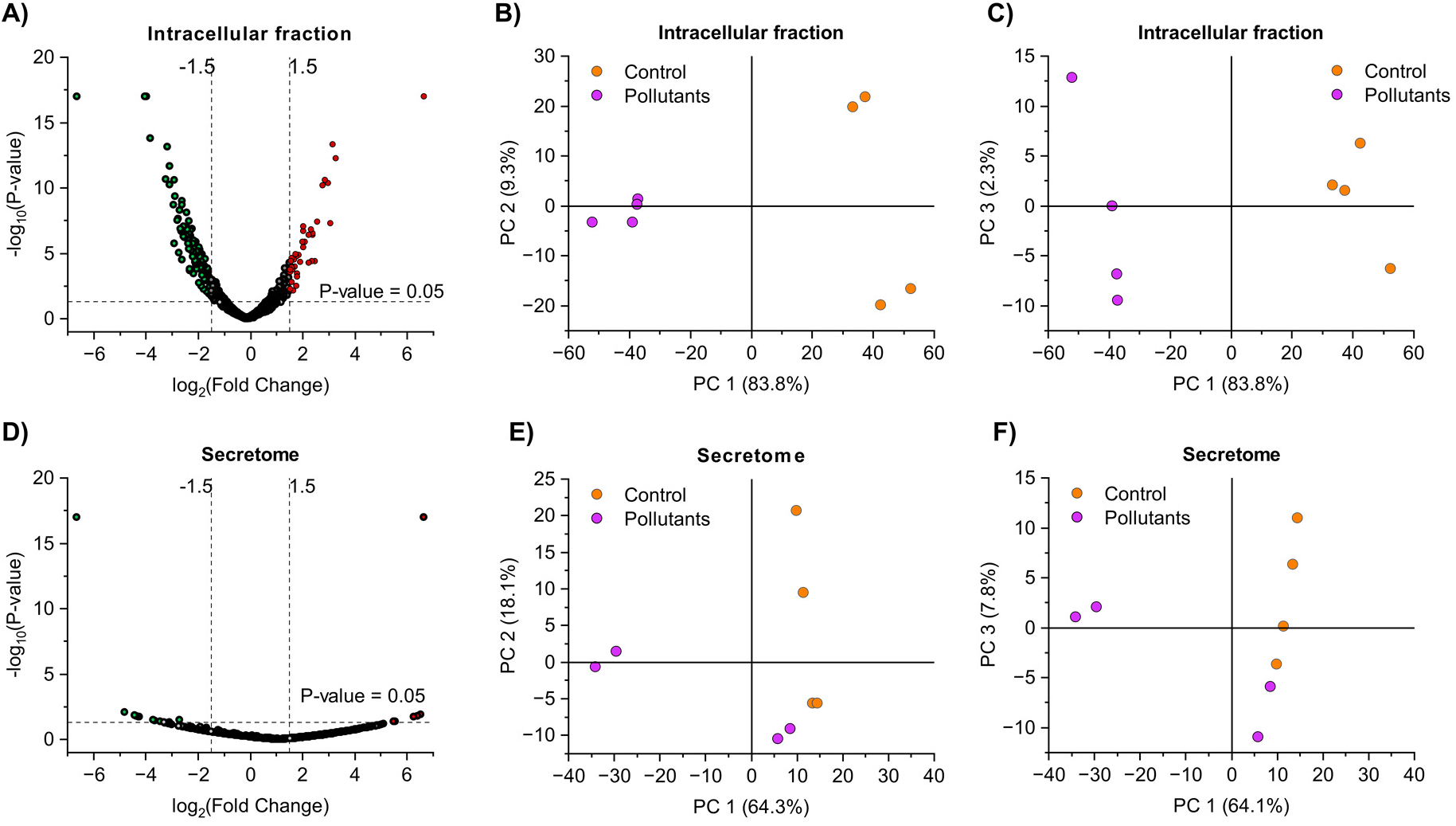
Proteomics analysis. A-C) Intracellular fraction. B-C) Secretome. A+D volcano plots demonstrating log2 fold changes. B+C and E+F Principal component (PC) analysis of proteomics data.

**FIG S7:**
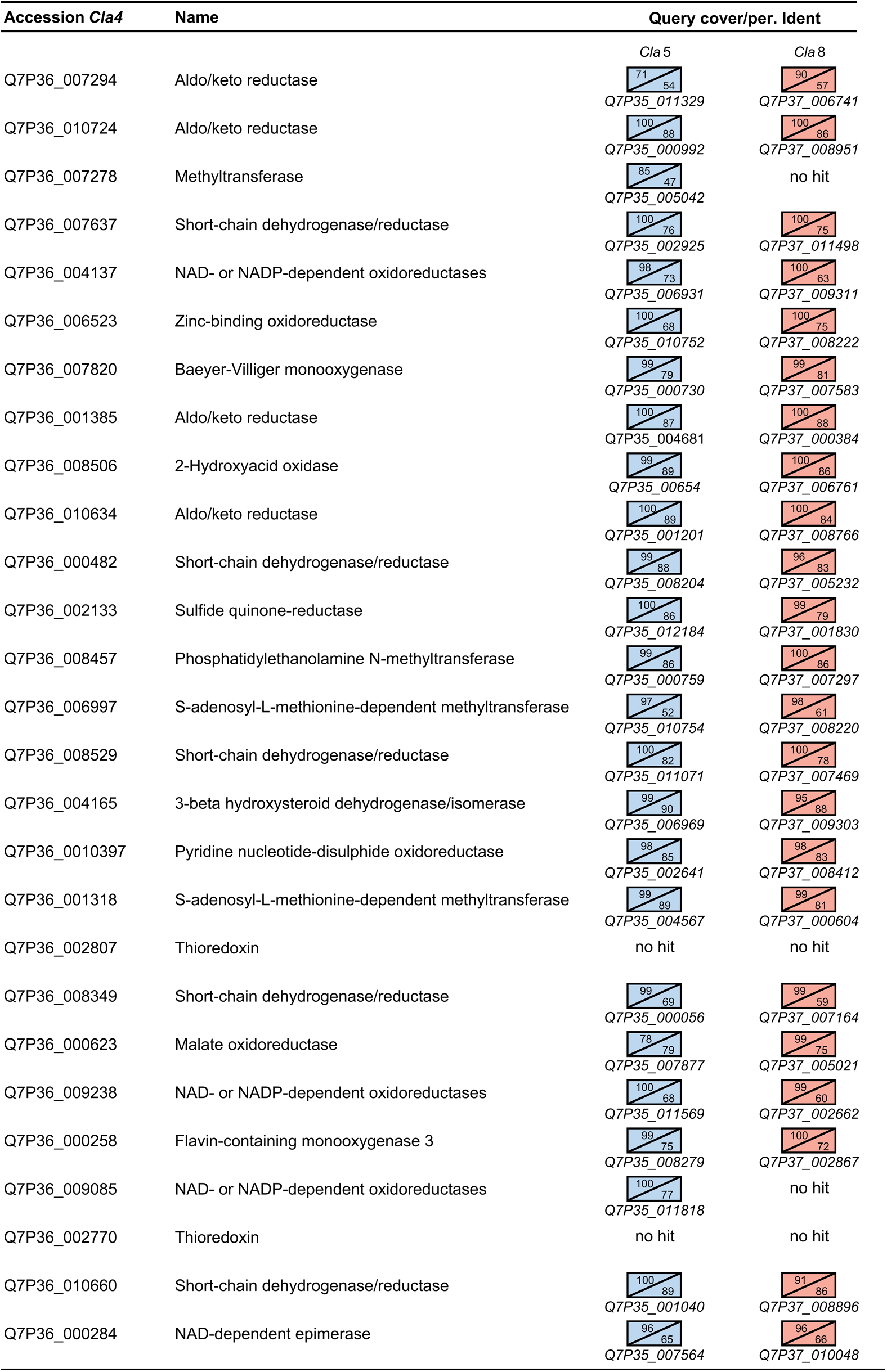
Conservation of upregulated oxidoreductases and methyltransferases within *Cla*5 and *Cla*8. Query coverage and percent identity are shown as percentages. Blue: *Cla*5 and red: *Cla*8. Accession numbers for homolog proteins are depicted under each box. No hit: Blast of a protein against the genomes of *Cla*5 or *Cla*8 gave no significant results.

## References

1. aus der Beek T, Weber F-A, Bergmann A, Hickmann S, Ebert I, Hein A, Küster A. 2016. Pharmaceuticals in the environment-Global occurrences and perspectives. Environ Toxicol Chem 35:823–835.

2. Li WC. 2014. Occurrence, sources, and fate of pharmaceuticals in aquatic environment and soil. Environ Pollut 187:193–201.

3. Fick J, Lindberg RH, Tysklind M, Larsson DGJ. 2010. Predicted critical environmental concentrations for 500 pharmaceuticals. Regul Toxicol Pharmacol 58:516–523.

4. Daughton CG. 2004. Non-regulated water contaminants: Emerging research. Environ Impact Assess Rev 24:711–732.

5. Fent K, Weston AA, Caminada D. 2006. Ecotoxicology of human pharmaceuticals. Aquat Toxicol 76:122–159.

6. Sanchez W, Sremski W, Piccini B, Palluel O, Maillot-Maréchal E, Betoulle S, Jaffal A, Aït-Aïssa S, Brion F, Thybaud E, Hinfray N, Porcher J-M. 2011. Adverse effects in wild fish living downstream from pharmaceutical manufacture discharges. Environ Int 37:1342–1348.

7. Nash JP, Kime DE, Van der Ven LTM, Wester PW, Brion F, Maack G, Stahlschmidt-Allner P, Tyler CR. 2004. Long-Term Exposure to Environmental Concentrations of the Pharmaceutical Ethynylestradiol Causes Reproductive Failure in Fish. Environ Health Perspect 112:1725–1733.

8. Lagesson A, Fahlman J, Brodin T, Fick J, Jonsson M, Byström P, Klaminder J. 2016. Bioaccumulation of five pharmaceuticals at multiple trophic levels in an aquatic food web - Insights from a field experiment. Sci Total Environ 568:208–215.

9. Patel M, Kumar R, Kishor K, Mlsna T, Pittman CU, Mohan D. 2019. Pharmaceuticals of emerging concern in aquatic systems: Chemistry, occurrence, effects, and removal methods. Chem Rev 119:3510–3673.

10. Kümmerer K, Dionysiou DD, Olsson O, Fatta-Kassinos D. 2019. Reducing aquatic micropollutants – Increasing the focus on input prevention and integrated emission management. Sci Total Environ 652:836–850.

11. Tijani JO, Fatoba OO, Madzivire G, Petrik LF. 2014. A Review of Combined Advanced Oxidation Technologies for the Removal of Organic Pollutants from Water. Water, Air, Soil Pollut 225:2102.

12. Magdeburg A, Stalter D, Schlüsener M, Ternes T, Oehlmann J. 2014. Evaluating the efficiency of advanced wastewater treatment: target analysis of organic contaminants and (geno-)toxicity assessment tell a different story. Water Res 50:35–47.

13. Mir-Tutusaus JA, Baccar R, Caminal G, Sarrà M. 2018. Can white-rot fungi be a real wastewater treatment alternative for organic micropollutants removal? A review. Water Res 138:137–151.

14. Cruz-Morató C, Lucas D, Llorca M, Rodriguez-Mozaz S, Gorga M, Petrovic M, Barceló D, Vicent T, Sarrà M, Marco-Urrea E. 2014. Hospital wastewater treatment by fungal bioreactor: Removal efficiency for pharmaceuticals and endocrine disruptor compounds. Sci Total Environ 493:365–376.

15. Lucas D, Badia-Fabregat M, Vicent T, Caminal G, Rodríguez-Mozaz S, Balcázar JL, Barceló D. 2016. Fungal treatment for the removal of antibiotics and antibiotic resistance genes in veterinary hospital wastewater. Chemosphere 152:301–308.

16. Cruz-Morató C, Ferrando-Climent L, Rodriguez-Mozaz S, Barceló D, Marco-Urrea E, Vicent T, Sarrà M. 2013. Degradation of pharmaceuticals in non-sterile urban wastewater by *Trametes versicolor* in a fluidized bed bioreactor. Water Res 47:5200–5210.

17. Tran NH, Urase T, Kusakabe O. 2010. Biodegradation Characteristics of Pharmaceutical Substances by Whole Fungal Culture Trametes versicolor and its Laccase. J Water Environ Technol 8:125–140.

18. Gallardo-Altamirano MJ, Maza-Márquez P, Montemurro N, Rodelas B, Osorio F, Pozo C. 2019. Linking microbial diversity and population dynamics to the removal efficiency of pharmaceutically active compounds (PhACs) in an anaerobic/anoxic/aerobic (A2O) system. Chemosphere 233:828–842.

19. Weber SD, Hofmann A, Pilhofer M, Wanner G, Agerer R, Ludwig W, Schleifer K-H, Fried J. 2009. The diversity of fungi in aerobic sewage granules assessed by 18S rRNA gene and ITS sequence analyses. FEMS Microbiol Ecol 68:246–254.

20. Maza-Márquez P, Vilchez-Vargas R, Kerckhof FM, Aranda E, González-López J, Rodelas B. 2016. Community structure, population dynamics and diversity of fungi in a full-scale membrane bioreactor (MBR) for urban wastewater treatment. Water Res 105:507–519.

21. González-Abradelo D, Pérez-Llano Y, Peidro-Guzmán H, Sánchez-Carbente M del R, Folch-Mallol JL, Aranda E, Vaidyanathan VK, Cabana H, Gunde-Cimerman N, Batista-García RA. 2019. First demonstration that ascomycetous halophilic fungi (*Aspergillus sydowii* and *Aspergillus destruens*) are useful in xenobiotic mycoremediation under high salinity conditions. Bioresour Technol 279:287–296.

22. Olicón-Hernández DR, Gómez-Silván C, Pozo C, Andersen GL, González-Lopez J, Aranda E. 2021. *Penicillium oxalicum* XD-3.1 removes pharmaceutical compounds from hospital wastewater and outcompetes native bacterial and fungal communities in fluidised batch bioreactors. Int Biodeterior Biodegradation 158:105179.

23. European Commission. 2022. Commission Implementing Decision (EU) 2022/1307 of 22 July 2022 establishing a watch list of substances for Union-wide monitoring in the field of water policy pursuant to Directive 2008/105/EC of the European Parliament and of the Council. Off J Eur Union.

24. Zhao D, Louise Leth M, Abou Hachem M, Aziz I, Jančič N, Luxbacher T, Hélix-Nielsen C, Zhang W. 2023. Facile fabrication of flexible ceramic nanofibrous membranes for enzyme immobilization and transformation of emerging pollutants. Chem Eng J 451:138902.

25. Widmer AF, Wiestner A, Frei R, Zimmerli W. 1991. Killing of nongrowing and adherent *Escherichia coli* determines drug efficacy in device-related infections. Antimicrob Agents Chemother 35:741–746.

26. Bensch K, Groenewald JZ, Meijer M, Dijksterhuis J, Jurjević Ž, Andersen B, Houbraken J, Crous PW, Samson RA. 2018. *Cladosporium* species in indoor environments. Stud Mycol 89:177–301.

27. Bensch K, Groenewald JZ, Braun U, Dijksterhuis J, de Jesús Yáñez-Morales M, Crous PW. 2015. Common but different: The expanding realm of *Cladosporium*. Stud Mycol 82:23–74.

28. Shin J, Kim J-E, Lee Y-W, Son H. 2018. Fungal Cytochrome P450s and the P450 Complement (CYPome) of *Fusarium graminearum*. Toxins (Basel) 10.3390/toxins10030112.

29. Drula E, Garron M-L, Dogan S, Lombard V, Henrissat B, Terrapon N. 2022. The carbohydrate-active enzyme database: functions and literature. Nucleic Acids Res 50:D571–D577.

30. Janusz G, Kucharzyk KH, Pawlik A, Staszczak M, Paszczynski AJ. 2013. Fungal laccase, manganese peroxidase and lignin peroxidase: Gene expression and regulation. Enzyme Microb Technol 52:1–12.

31. Berrin J-G, Rosso M-N, Abou Hachem M. 2017. Fungal secretomics to probe the biological functions of lytic polysaccharide monooxygenases. Carbohydr Res 448:155–160.

32. Ruiz-Herrera J, Ortiz-Castellanos L. 2019. Cell wall glucans of fungi. A review. Cell Surf 5:100022.

33. Rodríguez-Rodríguez CE, Jelić A, Llorca M, Farré M, Caminal G, Petrović M, Barceló D, Vicent T. 2011. Solid-phase treatment with the fungus *Trametes versicolor* substantially reduces pharmaceutical concentrations and toxicity from sewage sludge. Bioresour Technol 102:5602–5608.

34. Hultberg M, Ahrens L, Golovko O. 2020. Use of lignocellulosic substrate colonized by oyster mushroom (*Pleurotus ostreatus*) for removal of organic micropollutants from water. J Environ Manage 272:111087.

35. Cruz del Álamo A, Pariente MI, Martínez F, Molina R. 2020. *Trametes versicolor* immobilized on rotating biological contactors as alternative biological treatment for the removal of emerging concern micropollutants. Water Res 170:115313.

36. Hofmann U, Schlosser D. 2016. Biochemical and physicochemical processes contributing to the removal of endocrine-disrupting chemicals and pharmaceuticals by the aquatic ascomycete *Phoma* sp. UHH 5-1-03. Appl Microbiol Biotechnol 100:2381–2399.

37. Olicón-Hernández DR, Camacho-Morales RL, Pozo C, González-López J, Aranda E. 2019. Evaluation of diclofenac biodegradation by the ascomycete fungus *Penicillium oxalicum* at flask and bench bioreactor scales. Sci Total Environ 662:607–614.

38. Piyaviriyakul P, Boontanon N, Boontanon SK. 2021. Bioremoval and tolerance study of sulfamethoxazole using whole cell *Trichoderma harzianum* isolated from rotten tree bark. J Environ Sci Heal Part A 56:920–927.

39. Parshikov IA, Moody JD, Freeman JP, Lay Jr. JO, Williams AJ, Heinze TM, Sutherland JB. 2002. Formation of conjugates from ciprofloxacin and norfloxacin in cultures of *Trichoderma viride*. Mycologia 94:1–5.

40. Olicón-Hernández DR, Ortúzar M, Pozo C, González-López J, Aranda E. 2020. Metabolic Capability of *Penicillium oxalicum* to Transform High Concentrations of Anti-Inflammatory and Analgesic Drugs. Appl Sci 10.3390/app10072479.

41. Bensch K, Braun U, Groenewald JZ, Crous PW. 2012. The genus *Cladosporium*. Stud Mycol 72:1–401.

42. Ogórek R, Lejman A, Pusz W, Miłuch A, Miodyńska P. 2012. Characteristics and taxonomy of *Cladosporium* fungi. Mikol Lek 19(2):80– 85.

43. Alwatban MA, Hadi S, Moslem MA. 2014. Mycotoxin production in *Cladosporium* species influenced by temperature regimes. J Pure Appl Microbiol 8:4061–4069.

44. Lees-Haley PR. 2003. Toxic Mold and Mycotoxins in Neurotoxicity Cases: *Stachybotrys, Fusarium, Trichoderma, Aspergillus, Penicillium, Cladosporium, Alternaria, Trichothecenes*. Psychol Rep 93:561–584.

45. Pietsch C, Müller G, Mourabit S, Carnal S, Bandara K. 2020. Occurrence of Fungi and Fungal Toxins in Fish Feed during Storage. Toxins (Basel) 10.3390/toxins12030171.

46. Salvatore MM, Andolfi A, Nicoletti R. 2021. The Genus *Cladosporium*: A Rich Source of Diverse and Bioactive Natural Compounds. Molecules 10.3390/molecules26133959.

47. Mohamed GA, Ibrahim SRM. 2021. Untapped Potential of Marine-Associated *Cladosporium* Species: An Overview on Secondary Metabolites, Biotechnological Relevance, and Biological Activities. Mar Drugs. Department of Natural Products and Alternative Medicine, Faculty of Pharmacy, King Abdulaziz University, Jeddah 21589, Saudi Arabia. 10.3390/md19110645.

48. Viitanen H, Bjurman J. 1995. Mold growth on wood under fluctuating humidity conditions. Mater und Org 29:27–46.

49. Bland J, Gribble LA, Hamel MC, Wright JB, Moormann G, Bachand M, Wright G, Bachand GD. 2022. Evaluating changes in growth and pigmentation of *Cladosporium cladosporioides* and *Paecilomyces variotii* in response to gamma and ultraviolet irradiation. Sci Rep 12:12142.

50. Papazlatani C V, Kolovou M, Gkounou EE, Azis K, Mavriou Z, Testembasis S, Karaoglanidis GS, Ntougias S, Karpouzas DG. 2022. Isolation, characterization and industrial application of a *Cladosporium herbarum* fungal strain able to degrade the fungicide imazalil. Environ Pollut 301:119030.

51. Chen S, Liu C, Peng C, Liu H, Hu M, Zhong G. 2012. Biodegradation of Chlorpyrifos and Its Hydrolysis Product 3,5,6-Trichloro-2- Pyridinol by a New Fungal Strain *Cladosporium cladosporioides* Hu-01. PLoS One 7:e47205.

52. Birolli WG, de A. Santos D, Alvarenga N, Garcia ACFS, Romão LPC, Porto ALM. 2018. Biodegradation of anthracene and several PAHs by the marine-derived fungus *Cladosporium* sp. CBMAI 1237. Mar Pollut Bull 129:525–533.

53. Cortés-Espinosa D V, Fernández-Perrino FJ, Arana-Cuenca A, Esparza-García F, Loera O, Rodríguez-Vázquez R. 2006. Selection and Identification of Fungi Isolated from Sugarcane Bagasse and their Application for Phenanthrene Removal from Soil. J Environ Sci Heal Part A 41:475–486.

54. Llorca M, Badia-Fabregat M, Rodríguez-Mozaz S, Caminal G, Vicent T, Barceló D. 2017. Fungal treatment for the removal of endocrine disrupting compounds from reverse osmosis concentrate: Identification and monitoring of transformation products of benzotriazoles. Chemosphere 184:1054–1070.

55. Kózka B, Nałęcz-Jawecki G, Turło J, Giebułtowicz J. 2020. Application of *Pleurotus ostreatus* to efficient removal of selected antidepressants and immunosuppressant. J Environ Manage 273:111131.

56. Marco-Urrea E, Pérez-Trujillo M, Cruz-Morató C, Caminal G, Vicent T. 2010. Degradation of the drug sodium diclofenac by *Trametes versicolor* pellets and identification of some intermediates by NMR. J Hazard Mater 176:836–842.

57. Margot J, Maillard J, Rossi L, Barry DA, Holliger C. 2013. Influence of treatment conditions on the oxidation of micropollutants by *Trametes versicolor* laccase. N Biotechnol 30:803–813.

58. Nikolaivits E, Siaperas R, Agrafiotis A, Ouazzani J, Magoulas A, Gioti Α, Topakas E. 2021. Functional and transcriptomic investigation of laccase activity in the presence of PCB29 identifies two novel enzymes and the multicopper oxidase repertoire of a marine-derived fungus. Sci Total Environ 775:145818.

59. Aranda E, Godoy P, Reina R, Badia-Fabregat M, Rosell M, Marco-Urrea E, García-Romera I. 2017. Isolation of Ascomycota fungi with capability to transform PAHs: Insights into the biodegradation mechanisms of *Penicillium oxalicum*. Int Biodeterior Biodegradation 122:141–150.

60. Peidro-Guzmán H, Pérez-Llano Y, González-Abradelo D, Fernández-López MG, Dávila-Ramos S, Aranda E, Hernández DRO, García AO, Lira-Ruan V, Pliego OR, Santana MA, Schnabel D, Jiménez-Gómez I, Mouriño-Pérez RR, Aréchiga-Carvajal ET, del Rayo Sánchez- Carbente M, Folch-Mallol JL, Sánchez-Reyes A, Vaidyanathan VK, Cabana H, Gunde-Cimerman N, Batista-García RA. 2021. Transcriptomic analysis of polyaromatic hydrocarbon degradation by the halophilic fungus *Aspergillus sydowii* at hypersaline conditions. Environ Microbiol 23:3435–3459.

61. Lucero Camacho-Morales R, García-Fontana C, Fernández-Irigoyen J, Santamaría E, González-López J, Manzanera M, Aranda E. 2018. Anthracene drives sub-cellular proteome-wide alterations in the degradative system of *Penicillium oxalicum*. Ecotoxicol Environ Saf 159:127–135.

62. Feng M, Zhou J, Yu X, Mao W, Guo Y, Wang H. 2022. Insights into biodegradation mechanisms of triphenyl phosphate by a novel fungal isolate and its potential in bioremediation of contaminated river sediment. J Hazard Mater 424:127545.

63. Yu X, Mao C, Wang W, Kulshrestha S, Zhang P, Usman M, Zong S, Hilal MG, Fang Y, Han H, Li X. 2023. Reduction of metronidazole in municipal wastewater and protection of activated sludge system using a novel immobilized *Aspergillus tabacinus* LZ-M. Bioresour Technol 369:128509.

64. Marco-Urrea E, García-Romera I, Aranda E. 2015. Potential of non-ligninolytic fungi in bioremediation of chlorinated and polycyclic aromatic hydrocarbons. N Biotechnol 32:620–628.

65. Hoffmann F, Maser E. 2007. Carbonyl Reductases and Pluripotent Hydroxysteroid Dehydrogenases of the Short-chain Dehydrogenase/reductase Superfamily. Drug Metab Rev 39:87–144.

66. Novotna R, Wsol V, Xiong G, Maser E. 2008. Inactivation of the anticancer drugs doxorubicin and oracin by aldo–keto reductase (AKR) 1C3. Toxicol Lett 181:1–6.

67. Penning TM. 2015. The aldo-keto reductases (AKRs): Overview. Chem Biol Interact 234:236–246.

68. Krueger SK, Williams DE. 2005. Mammalian flavin-containing monooxygenases: structure/function, genetic polymorphisms and role in drug metabolism. Pharmacol Ther 106:357–387.

69. Lennard L. 2010. Methyltransferases. Compr Toxicol Second Ed 4:435–457.

70. Soares PRS, Birolli WG, Ferreira IM, Porto ALM. 2021. Biodegradation pathway of the organophosphate pesticides chlorpyrifos, methyl parathion and profenofos by the marine-derived fungus *Aspergillus sydowii* CBMAI 935 and its potential for methylation reactions of phenolic compounds. Mar Pollut Bull 166:112185.

71. Juan CA, Pérez de la Lastra JM, Plou FJ, Pérez-Lebeña E. 2021. The Chemistry of Reactive Oxygen Species (ROS) Revisited: Outlining Their Role in Biological Macromolecules (DNA, Lipids and Proteins) and Induced Pathologies. Int J Mol Sci 10.3390/ijms22094642.

72. Holmgren A, Lu J. 2010. Thioredoxin and thioredoxin reductase: Current research with special reference to human disease. Biochem Biophys Res Commun 396:120–124.

73. Collet J-F, Messens J. 2010. Structure, Function, and Mechanism of Thioredoxin Proteins. Antioxid Redox Signal 13:1205–1216.

74. Pluskal T, Castillo S, Villar-Briones A, Orešič M. 2010. MZmine 2: Modular framework for processing, visualizing, and analyzing mass spectrometry-based molecular profile data. BMC Bioinformatics 11:395.

75. Wang M, Carver JJ, Phelan V V, Sanchez LM, Garg N, Peng Y, Nguyen DD, Watrous J, Kapono CA, Luzzatto-Knaan T, Porto C, Bouslimani A, Melnik A V, Meehan MJ, Liu W-T, Crüsemann M, Boudreau PD, Esquenazi E, Sandoval-Calderón M, Kersten RD, Pace LA, Quinn RA, Duncan KR, Hsu C-C, Floros DJ, Gavilan RG, Kleigrewe K, Northen T, Dutton RJ, Parrot D, Carlson EE, Aigle B, Michelsen CF, Jelsbak L, Sohlenkamp C, Pevzner P, Edlund A, McLean J, Piel J, Murphy BT, Gerwick L, Liaw C-C, Yang Y-L, Humpf H-U, Maansson M, Keyzers RA, Sims AC, Johnson AR, Sidebottom AM, Sedio BE, Klitgaard A, Larson CB, Boya P CA, Torres-Mendoza D, Gonzalez DJ, Silva DB, Marques LM, Demarque DP, Pociute E, O’Neill EC, Briand E, Helfrich EJN, Granatosky EA, Glukhov E, Ryffel F, Houson H, Mohimani H, Kharbush JJ, Zeng Y, Vorholt JA, Kurita KL, Charusanti P, McPhail KL, Nielsen KF, Vuong L, Elfeki M, Traxler MF, Engene N, Koyama N, Vining OB, Baric R, Silva RR, Mascuch SJ, Tomasi S, Jenkins S, Macherla V, Hoffman T, Agarwal V, Williams PG, Dai J, Neupane R, Gurr J, Rodríguez AMC, Lamsa A, Zhang C, Dorrestein K, Duggan BM, Almaliti J, Allard P-M, Phapale P, Nothias L-F, Alexandrov T, Litaudon M, Wolfender J-L, Kyle JE, Metz TO, Peryea T, Nguyen D-T, VanLeer D, Shinn P, Jadhav A, Müller R, Waters KM, Shi W, Liu X, Zhang L, Knight R, Jensen PR, Palsson BØ, Pogliano K, Linington RG, Gutiérrez M, Lopes NP, Gerwick WH, Moore BS, Dorrestein PC, Bandeira N. 2016. Sharing and community curation of mass spectrometry data with Global Natural Products Social Molecular Networking. Nat Biotechnol 34:828–837.

76. Shannon P, Markiel A, Ozier O, Baliga NS, Wang JT, Ramage D, Amin N, Schwikowski B, Ideker T. 2003. Cytoscape: A Software Environment for Integrated Models of Biomolecular Interaction Networks. Genome Res 13:2498–2504.

77. Petersen C, Sørensen T, Westphal KR, Fechete LI, Sondergaard TE, Sørensen JL, Nielsen KL. 2022. High molecular weight DNA extraction methods lead to high quality filamentous ascomycete fungal genome assemblies using Oxford Nanopore sequencing. Microb Genomics 8:000816.

78. Li H. 2018. Minimap2: pairwise alignment for nucleotide sequences. Bioinformatics 34:3094–3100.

79. Li H. 2016. Minimap and miniasm: fast mapping and de novo assembly for noisy long sequences. Bioinformatics 32:2103–2110.

80. Vaser R, Sović, I, Sović, S, Nagarajan N, Šikić1 M, Šikić1 Š. 2017. Fast and accurate de novo genome assembly from long uncorrected reads 10.1101/gr.214270.116.

81. Seppey M, Manni M, Zdobnov EM. 2019. BUSCO: Assessing Genome Assembly and Annotation Completeness BT - Gene Prediction: Methods and Protocols. Methods Mol Biol 1962:227–245.

82. Stanke M, Steinkamp R, Waack S, Morgenstern B. 2004. AUGUSTUS: a web server for gene finding in eukaryotes. Nucleic Acids Res 32:W309–W312.

83. Cantalapiedra CP, Hernández-Plaza A, Letunic I, Bork P, Huerta-Cepas J. 2021. eggNOG-mapper v2: Functional Annotation, Orthology Assignments, and Domain Prediction at the Metagenomic Scale. Mol Biol Evol 38:5825–5829.

84. Paysan-Lafosse T, Blum M, Chuguransky S, Grego T, Pinto BL, Salazar GA, Bileschi ML, Bork P, Bridge A, Colwell L, Gough J, Haft DH, Letunić I, Marchler-Bauer A, Mi H, Natale DA, Orengo CA, Pandurangan AP, Rivoire C, Sigrist CJA, Sillitoe I, Thanki N, Thomas PD, Tosatto SCE, Wu CH, Bateman A. 2023. InterPro in 2022. Nucleic Acids Res 51:D418–D427.

85. White TJ, Bruns T, Lee S, Taylor J. 1990. Amplification and direct sequencing of fungal ribosomal RNA genes for phylogenetics. PCR Protoc a Guid to methods Appl 18:315–322.

86. Carbone I, Kohn LM. 1999. A method for designing primer sets for speciation studies in filamentous ascomycetes. Mycologia 91:553– 556.

87. Yin Y, Mao X, Yang J, Chen X, Mao F, Xu Y. 2012. DbCAN: A web resource for automated carbohydrate-active enzyme annotation. Nucleic Acids Res 40:445–451.

88. Jain C, Rodriguez-R LM, Phillippy AM, Konstantinidis KT, Aluru S. 2018. High throughput ANI analysis of 90K prokaryotic genomes reveals clear species boundaries. Nat Commun 9:5114.

89. Teufel F, Almagro Armenteros JJ, Johansen AR, Gíslason MH, Pihl SI, Tsirigos KD, Winther O, Brunak S, von Heijne G, Nielsen H. 2022. SignalP 6.0 predicts all five types of signal peptides using protein language models. Nat Biotechnol 40:1023–1025.

90. Hallgren J, Tsirigos KD, Pedersen MD, Almagro Armenteros JJ, Marcatili P, Nielsen H, Krogh A, Winther O. 2022. DeepTMHMM predicts alpha and beta transmembrane proteins using deep neural networks. bioRxiv 2022.04.08.487609.

91. Thumuluri V, Almagro Armenteros JJ, Johansen AR, Nielsen H, Winther O. 2022. DeepLoc 2.0: multi-label subcellular localization prediction using protein language models. Nucleic Acids Res 50:W228–W234.

92. Altschul SF, Gish W, Miller W, Myers EW, Lipman DJ. 1990. Basic local alignment search tool. J Mol Biol 215:403–410.

